# Genomic footprints of repeated evolution of CAM photosynthesis in tillandsioid bromeliads

**DOI:** 10.1101/495812

**Authors:** Marylaure De La Harpe, Margot Paris, Jaqueline Hess, Michael H. J. Barfuss, Martha L. Serrano-Serrano, Arindam Ghatak, Palak Chaturvedi, Wolfram Weckwerth, Walter Till, Nicolas Salamin, Ching Man Wai, Ray Ming, Christian Lexer

## Abstract

The adaptive radiation of Bromeliaceae (pineapple family) is one of the most diverse among Neotropical flowering plants. Diversification in this group was facilitated by several ‘key innovations’ including the transition from C3 to CAM photosynthesis. We used a phylogenomic approach complemented by differential gene expression (RNA-seq) and targeted metabolite profiling to address the patterns and mechanisms of C3/CAM evolution in the extremely species-rich bromeliad genus Tillandsia and related taxa. Evolutionary analyses at a range of different levels (selection on protein-coding genes, gene duplication and loss, regulatory evolution) revealed three common themes driving the evolution of CAM: response to heat and drought, alterations to basic carbohydrate metabolism, and regulation of organic acid storage. At the level of genes and their products, CAM/C3 shifts were accompanied by gene expansion of a circadian regulator, re-programming of ABA-related gene expression, and adaptive sequence evolution of an enolase, effectively linking carbohydrate metabolism to ABA-mediated stress response. These changes include several pleiotropic regulators, which facilitated the evolution of correlated adaptive traits during a textbook adaptive radiation.

---

Species radiations have traditionally been studied primarily from a macro-evolutionary perspective within a phylogenetic framework^1–3^. The recent “-omics” revolution and novel analytical tools increasingly allow evolutionary biologists to address the dynamics of genomic variation across time scales^4–7^. This potentially allows evolutionary geneticists to identify the major sources of genetic variation that fuel evolutionary radiations^8^.

The Bromeliaceae family (>3000 species) represents a ‘textbook adaptive radiation’ in flowering plants^9^. Numerous adaptive traits or ‘key innovations’ vary among species of this large Neotropical family, including the epiphytic growth habit (life on trees), presence or absence of water-impounding leaves (tank-forming rosettes), absorptive trichomes, leaf succulence, and Crassulacean Acid Metabolism (CAM) photosynthesis^9–11^. Chromosome counts in the family point to predominant diploidy with a remarkably widespread 2n=50 or 2n=48 chromosomes and a comparatively compact range of DNA content from 0.85 to 2.23pg/2C, suggesting a largely homoploid radiation^12^.

The highly species-rich genus *Tillandsia* L. (ca. 650 species) of the Tillandsioidae subfamily of Bromeliaceae exhibits great variation in life habits (epiphytic, terrestrial, and rock-growing), photosynthetic pathways (C3 and CAM), pollination syndromes (birds and insects), the presence or absence of absorptive trichomes and leaf succulence, and several other adaptive phenotypic traits^9–11,13,14^. These trait differences appear to have evolved in a correlated, contingent manner, giving rise to adaptive syndromes of correlated characters^11,15^. Most obviously, these trait associations manifest themselves in so-called “green” *Tillandsia* phenotypes and species adapted to cool, moist habitats, typically exhibiting C3 photosynthesis, neither pronounced absorptive trichome cover nor succulence, and widespread formation of tank rosettes. This contrasts with so-called “grey” *Tillandsia* phenotypes and forms with a strong tendency to express CAM photosynthesis, dense absorptive trichome cover and pronounced succulence, and strong association with warm, highly irradiated habitats in regions with low rainfall^9,11^. Although shifts in these adaptive traits have long been hypothesized to be drivers of adaptive radiation in this group^9,11,14^, little to nothing is known about their genetic basis, with the notable exception of CAM photosynthesis.

The CAM pathway, known from at least 35 different plant families, represents an adaptation for increased water use efficiency by shifting CO2 assimilation to the night time, thus allowing stomata to be closed during the day and thereby reducing water loss^16,17^. CAM entails complex diel patterns of gene expression, post-translational regulation, and metabolic fluxes that are only starting to be understood at a whole systems level^18^.

Here, we combine a range of experimental approaches to shed light on the genetic basis and evolution of CAM photosynthesis and correlated traits in the adaptive radiation of tillandsioid bromeliads. To this end, we studied 28 accessions of Tillandsioideae including 25 species of *Tillandsia* sensu lato, capturing much of the available variation in the CAM adaptive syndrome. We first used whole genome phylogenetics to ask whether repeated C3/CAM trait shifts in this radiation are more likely to have arisen independently or by wide-spread gene flow. Next, we examined genome-wide signatures of branch-specific selection and gene duplication/loss accompanying CAM-related trait shifts. We then zoomed in on time-dependent metabolic and transcriptomic changes in representative species. We show that repeated xeric (heat/drought) adaptation including CAM involves not only structural genes, but also highly pleiotropic regulators.

## Results

### Phenotypic variation captured

A total of 28 accessions representing 25 species of *Tillandsia* and closely related genera were studied to achieve representative sampling of phenotypic variation from the adaptive radiation of this group^9,14^, including also two outgroup taxa, *Alcantarea trepida* and *Vriesea itatiaiae*. We aimed to represent the range of photosynthetic syndromes ranging from typical C3 to typical CAM^10,19^. The CAM species sampled for this study display several phenotypic traits thought to represent adaptations to xeric conditions^9^ that likely evolved in a correlated, contingent manner^11^. Notwithstanding taxon-specific idiosyncrasies, there is a broad association between CAM, increased succulence, reduced leaf evaporation properties, dense trichomes, and the absence of a water tank in CAM tillandsioids (**Fig. 1**)^11^.

**Figure 1.**
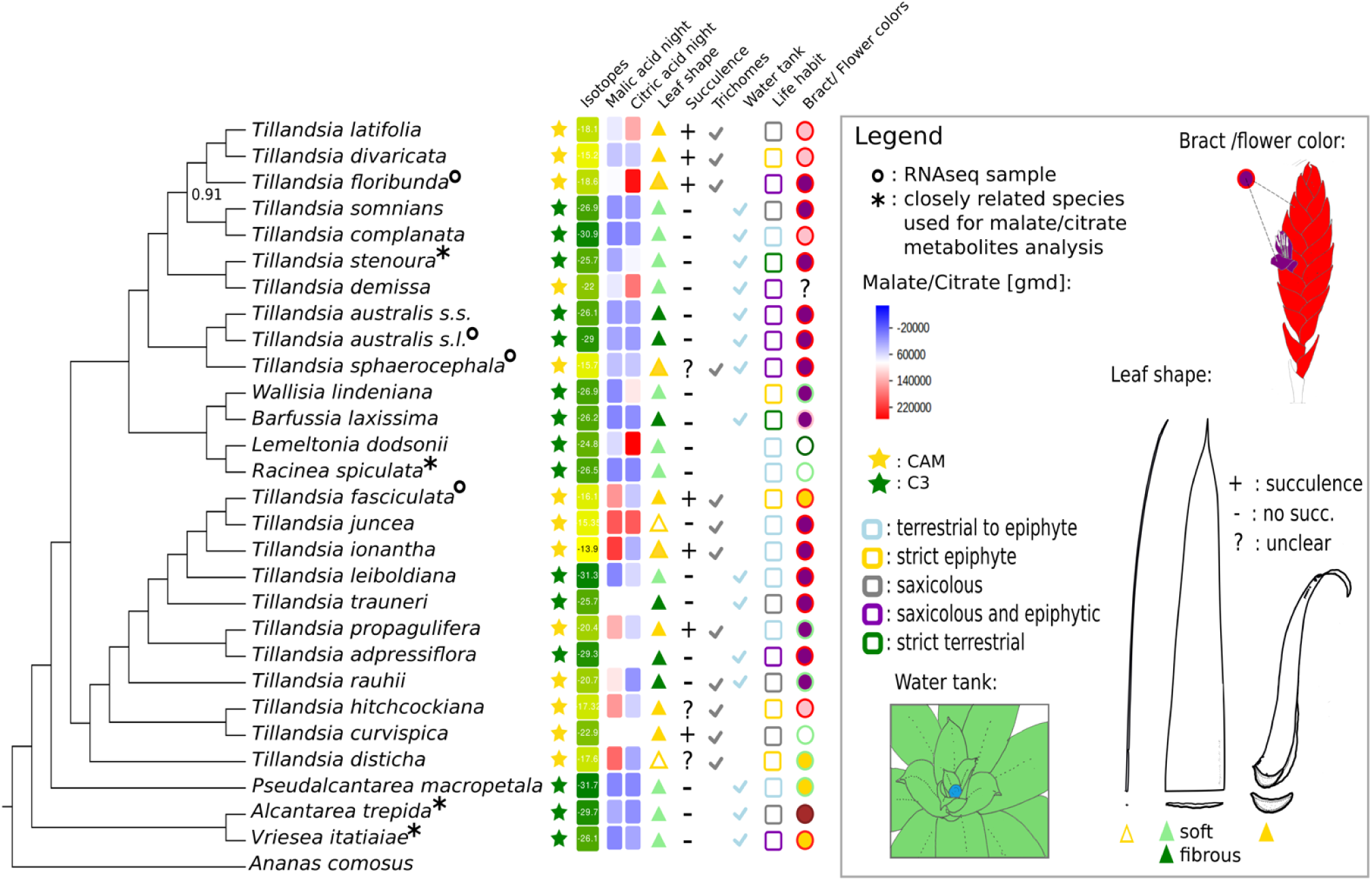
Coalescent-based phylogeny of 28 whole-genome sequenced accessions for 25 tillandsioid bromeliad species and *Ananas comosus* outgroup. Annotations display grouping into C3 (green stars) and CAM (yellow stars) species according to carbon isotope (δ^13^C) phenotypes. Measured carbon isotope ratios, night-time malate and citrate concentrations from targeted metabolite analysis, and differences in six other putatively adaptive plant traits are indicated in the legend box on the right. Samples used for RNA-seq expression profiling are indicated by a circle, samples with an asterisk represent species for which a close relative was used in the metabolite analysis.

### Phylogenomic relationships among C3 and CAM taxa

Whole genome sequencing of all species (**SI Table 1**) yielded 5,646,174 high-quality SNPs with an average coverage of 17.7x (median coverage: 20.6x). Coalescent-based reconstruction using ASTRAL resulted in a well resolved tree and we recovered major clades identified in previous molecular systematic work^14^. The coalescent tree was largely congruent with a maximum likelihood phylogeny estimated using RAxML (**SI Fig. 1, SI Text**). Based on the traditional sorting of taxa into discrete C3 and CAM phenotypes, our phylogenetic tree suggests a minimum of five independent transitions between C3 and CAM photosynthesis among the species sampled (**Fig. 1**). Placement of CAM and C3 taxa in the SplitsTree^20^ network indicate no apparent signals of evolutionary reticulation at this phylogenetic scale (SI **Fig. 2**).

### Molecular phenotypes capturing photosynthetic syndromes

Carbon isotope ratios recovered under greenhouse conditions indicate a continuum of values ranging from typical C3 to fairly strong CAM (**Fig. 1**), following commonly used thresholds^10,16^. Many species in our sample set displayed typical C3 carbon isotope (δ^13^C) phenotypes far beyond −20‰ and in fact reaching as far into the C3 extreme as −30‰ (labelled green in **Fig. 1**). On the other end of the C3/CAM continuum, *T. ionantha* exhibited a δ^13^C value of only −13.9‰, indicating it represents a so-called ‘strong’ CAM species (labelled in yellow in **Fig. 1**). Species may vary in their degree of night time carbon utilization, in particular those with drought-inducible, facultative CAM phenotypes^16,21^. Since carbon isotope measurements were taken under standardized, well-watered conditions, it is thus possible that our phenotyping classified some inducible CAM species as C3. Other phenotyping methods such as acidity under drought conditions^16,21^ were not practicable for our experimental set-up involving highly divergent phenotypes from precious living collections, which required careful consideration to avoid loss of unique accessions (**Supplemental Information**). In effect, we used isotopic ratios as a proxy to partition species according to the extremes of the CAM/C3 distribution for evolutionary analyses. This is a conservative and pragmatic strategy, since phenotyping error would likely diminish the signal-to-noise ratio.

We surveyed metabolic phenotypes of all species included in this study using GC-TOF-MS for metabolomic profiling of green tissues sampled at 11am and 1am, congruent with our sampling also used for gene expression profiling (below). Species also used for RNA-seq were represented by up to three biological replicates. Partial Least Squares Discriminant Analysis (PLS-DA) of 32 putatively identified metabolic compounds (comprising mainly amino acids, carbohydrates and organic acids; **SI Table 2**) indicated broad metabolic differentiation between CAM and C3 plants, especially along the second principal axis for both sampling time points, day and night (**Fig. 2**). These patterns were driven primarily by organic acids such as malic and fumaric acid as expected for CAM plants^9,22^, but also by soluble sugars. Closer inspection of organic acids revealed complex patterns of compound accumulation across taxa and sampling time points (**Fig. 1; Table S2**). Most conspicuously, species with strong CAM phenotypes in our study such as *T. ionantha* and *T. fasciculata* showed strongly increased night-time malic acid abundances, compared to most C3 species studied (**Fig. 1**). In some of the weaker CAM species such as *T. floribunda*, night-time malic acid abundances were inconspicuous, but citric acid abundances were increased instead. *Tillandsia australis*, our C3 reference species for expression profiling, did not exhibit increased accumulation of either CAM-related carbohydrate. A general pattern emerged with respect to carbohydrates (**Fig. 2C**), with more abundant soluble sugars in C3 compared to CAM plants (**Fig. 2C**).

**Figure 2.**
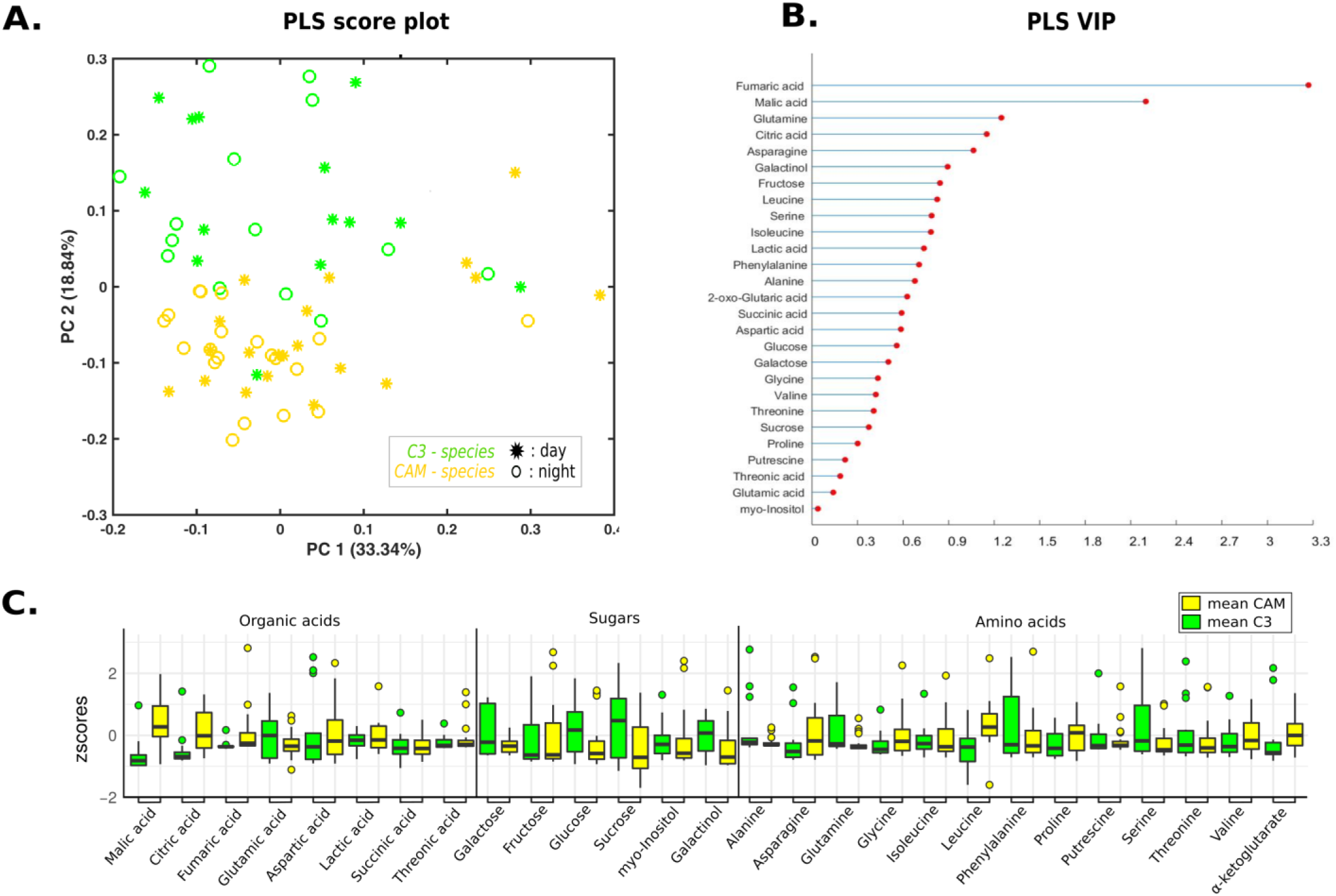
Partial Least Squares - Discriminant Analysis (PLS-DA) of targeted metabolite data. GC-TOF-MS analysis of green tissue sampled during the day (11 AM) and night (1 AM). **A**, PLS-DA score plot, C3/CAM day and night affiliation indicated in legend. **B**, Loadings. **C**, Boxplots of photosynthesis-relevant metabolites measured in C3 and CAM plants at 1AM. The y-scale was truncated at a z-score of 3 for better visibility of interquartile ranges (boxes) and medians (solid lines).

### Branch-specific positive selection in coding sequences

To identify genes that underwent adaptive protein evolution during C3/CAM transitions, we scanned the gene space of all genome-sequenced species for branch-specific signatures of positive selection^23^. Stringently implemented tests for positive selection in the coding regions of 13,603 genes revealed 22 genes that have apparently undergone adaptive protein evolution along branches relevant to C3/CAM shifts (**SI Table 3**). This includes two transcription factors (TFs) and eight genes of relevance in the context of the xeric adaptive syndrome associated with CAM in tillandsioids (**Fig. 1**). Eleven of the 22 genes under selection were identified as being differentially expressed in one or more of our CAM-related temporal and interspecific DE comparisons (**SI Table 3**; below). The 22 selected genes also include two genes involved in regulation of carbohydrate fluxes in plant tissues, an enolase (gene Aco020962; **SI Figs 6 and 7**) and a glucose6phosphate dehydrogenase (G6PD; gene Aco012435). The former encodes a homolog of AtENO2/LOS2, a bifunctional gene encoding both an enolase that catalyses the conversion of 2-phosphoglycerate to phosphoenolpyruvate, and MBP-1, a transcriptional repressor that plays a role in abscisic acid (ABA)-mediated response to abiotic stress^24^.

### Gene family evolution

We used birth-death models implemented in CAFÉ^25^ to investigate the impact of gene duplications and losses on the repeated evolution of the CAM correlated trait syndrome. Copy number variant (CNV) calling in the subset of species sequenced with high coverage (**Fig. 3**) identified gains in 2,808 and losses in 1,749 genes out of the 19,728 genes surveyed for CNVs. Clustering of the *A. comosus* proteome into gene families resulted in a total of 8,418 clusters of which 7,216 had non-zero counts in at least one *Tillandsia* species and were retained for further analysis. Estimation of gene birth (*λ*) and death (μ) rates revealed distinct evolutionary dynamics among different clades within *Tillandsia* (**Fig. 3**). Rates of duplication and loss were almost threefold higher in the clade containing species of the subg. *Tillandsia* with a Central and North American distribution, including *T. fasciculata*, *T. juncea*, *T. leiboldiana* and *T. trauneri****^26^*** when compared to the rest of the tree (λ_2_ = 0.002841, μ_2_ = 0.000865 and λ_1_ = 0.000795, μ_1_ = 0.000239, respectively). Mining for gene families with an increase in duplication or loss rate in association with trait shifts (C3 to CAM) along the tree resulted in five candidate gene families showing rate increases along more than one branch (**Table 1**). At least two of these are directly relevant to the adaptive trait syndrome associated with CAM plants: Cluster_1286 consists of a family of galactinol synthases and duplications in this family are all found in Aco001744, a homolog of GolS1 in *Arabidopsis thaliana*. In *A. thaliana*, this protein is involved in the production of raffinose, an osmoprotectant, and expressed under the control of heat shock factors in response to a combination of high light and heat stress^27^. The metabolite data also point to this metabolic checkpoint of adaptation, galactinol being a metabolite marker discriminating C3 and CAM plants (**Fig. 2C**). Cluster_7372, on the other hand, contains a single gene, Aco019534, which encodes a homolog of XAP5 CIRCARDIAN TIMEKEEPER (XCT) in *A. thaliana* (**Fig. 3**). AtXCT is a regulatory protein involved in regulation of circadian period length, developmental processes such as tissue greening and chloroplast development and important for sugar- and light quality-dependent ethylene signalling related to growth^28,29^. Moreover, AtXCT is known to be a key player in regulation of all three major classes of small RNAs in *A. thaliana*^30^.

**Table 1.** Gene families with putative association to CAM/C3 correlated trait syndrome

| Cluster | F | P-value<br>(FDR<br>adj.) | Annotation | Direction |
| --- | --- | --- | --- | --- |
| cluster_759 | 5.7640 | 0.0000 | Ypt/Rab-GAP domain of gyp1p superfamily protein | Loss |
| cluster_1286 | 4.6867 | 0.0000 | Galactinol synthase 1 | Gain |
| cluster_7372 | 4.4536 | 0.0000 | XAP5 CIRCADIAN TIMEKEEPER | Gain |
| cluster_392 | 3.4649 | 0.0000 | Thioredoxin | Gain |
| cluster_2314 | 6.3453 | 0.0243 | F-box family protein | Loss |
| cluster_1521 | 5.6398 | 0.0483 | TCP family transcription factor | Gain |

**Figure 3.**
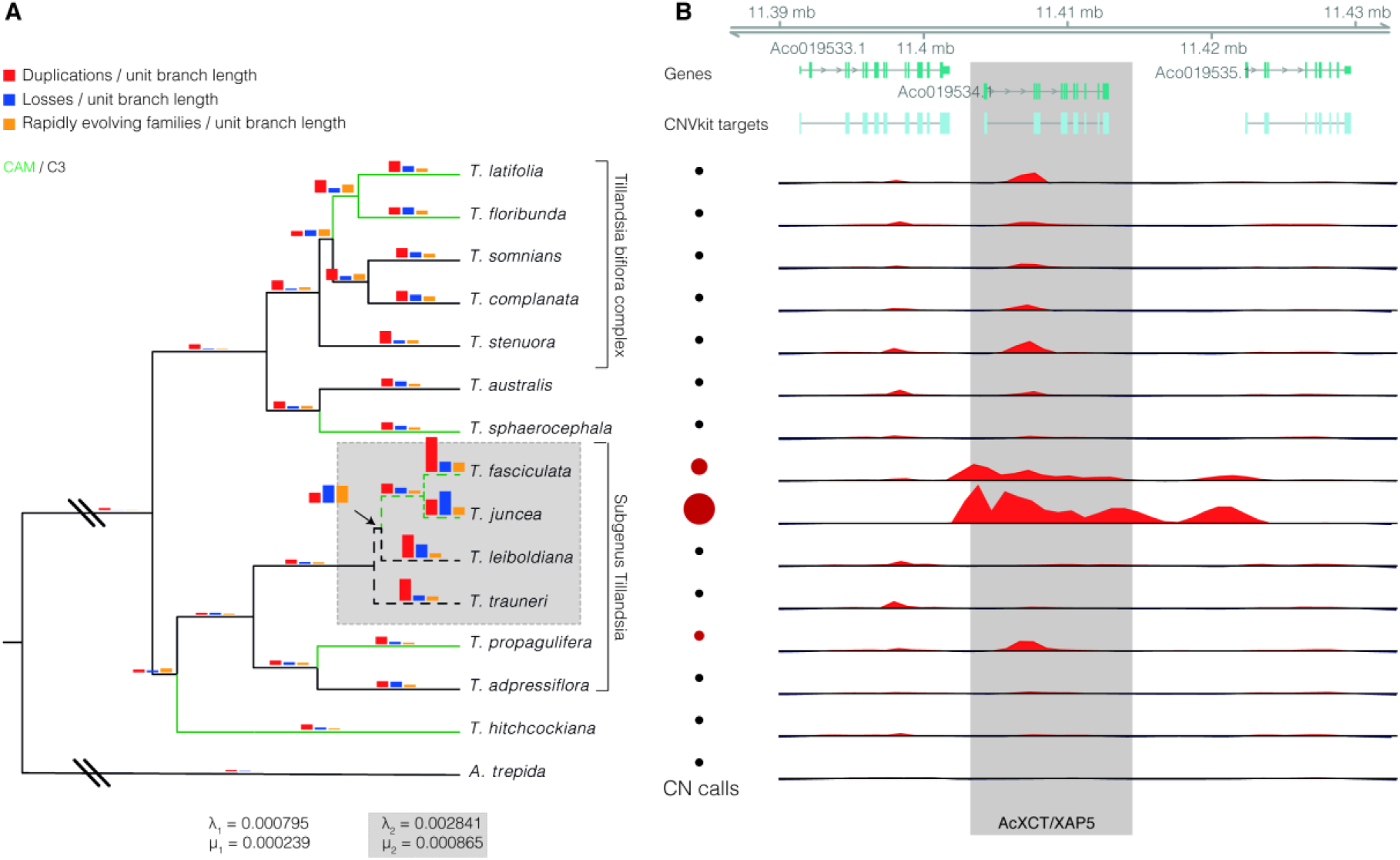
High-coverage samples included in analysis of copy number (CN) variants. Bar plots indicate inferred numbers of duplications (red) and losses (blue) by CAFÉ^24^ and numbers of families evolving at significantly different rates (P<0.01) compared to the genomic background (orange), all scaled by the lengths of the respective branches. Genome-wide rates of duplication (λ) and loss (μ) are given for subgenus *Tillandsia* (λ*_2_*, μ_2_, grey shading) and the rest of tree (λ_1_, μ_1_). Coloring on branches indicates whether branches are CAM (green) or C3 (black). The right panel illustrates estimated CNs and coverage distributions for AtXCT homolog Aco019534, one of the six gene families whose rates of duplication and loss showed significant association with the CAM phenotype (Table 1).

### Transcriptome-wide gene expression profiling across the C3/CAM continuum

To identify genes potentially involved in C3/CAM transitions and related trait shifts, we examined transcriptome-wide changes in gene expression by RNA sequencing (RNA-seq). One C3 species (*Tillandsia australis*; *Taust* from here onwards) and three CAM species (*T. sphaerocephala, Tspha; T. fasciculata, Tfasc; T. floribunda, Tflor*) were selected for RNA-seq (**Fig. 1**; **SI Table 1**). These species were chosen based on their range of carbon isotope phenotypes, the local availability of biological replicates, and because they represent CAM/C3 comparisons with different evolutionary distances between each CAM species and the C3 reference taxon (**Fig. 1**). *Taust*, our C3 reference taxon exhibited δ^13^C values of −26 to −29‰, clearly beyond the −20‰ threshold commonly used to classify C3 plants^16^. This species also exhibited all other phenotypic features expected for C3 bromeliads, including tank-forming rosettes, no succulence, and absence of dense trichome cover. The three CAM taxa sampled for expression profiling exhibited a broad range of CAM-like δ^13^C values and morphological features (**Supplemental Information**). Species were sampled at 1am and 11am, corresponding to the highest and lowest peaks, respectively of net CO_2_ assimilation rates (phase I, carboxylation at night and phase III, decarboxylation at day) and apparent PEPC kinase (PPCK) activation state of the CAM species *T. usneoides^31^*. Between 18,439 and 20,378 genes were successfully recovered per individual sample, representing between 68.2% and 75.4% of the 27,024 annotated genes presented in the *A. comosus* reference genome assembly. Multi-dimensional scaling (MDS) of RNA-seq data revealed clear clustering of biological replicates within each species, and patterns of interspecific differentiation along the first (=horizontal) axis broadly reflected the known phylogenetic relationships among the studied taxa (SI **Fig. 3**).

#### Intraspecific day/night comparisons

Comparisons of our day and night sampling time points within each species revealed that between 82 (*Tflor*) and 1354 (*Tfasc*) genes were differentially expressed at FDR 5% and log fold change (LFC) >1 between day and night conditions across the four tested species (**SI Fig. 4A**). On average, 65.4% of them were over-expressed during the night, with few apparent differences shared among the three CAM species to the exclusion of our C3 reference taxon (**SI Fig. 4A**). Only two genes showed diurnal regulation in all three CAM but not the C3 species. One of these, Aco003903 encodes a regulatory protein homologous to *A. thaliana* ABF2/ABF3. In *A. thaliana*, this gene family is involved in abscisic acid (ABA)-mediated stress response to sugar-, salt-, and osmotic stress^32^. The other gene (Aco016050) is of unknown function but encodes an ACT amino acid binding domain (PF01842) and may therefore play a role in amino acid metabolism. Differential gene expression (DE) results and gene ontology (GO) enrichments yielded patterns of up- and down-regulated genes and expressed metabolic pathways consistent with expected differences between day and night (SI **Fig. 5; SI Text**), including a range of photosynthesis-related GO terms.

#### Interspecific C3/CAM comparisons

Comparisons between each of our CAM species and our C3 reference taxon revealed between 1302 (*Tflor*) and 2757 (*Tfasc*) genes that were DE between CAM and C3 species during day conditions (11AM) with 49.9% of genes significantly up-regulated in CAM species. Under night conditions (1AM), 51.7% of the genes were up-regulated in CAM species, affecting between 1460 ***(Tflor)*** and 3110 ***(Tfasc)*** genes (**Fig. 4A**). Hundreds of DE transcripts were shared among all three CAM species in these comparisons, with many more shared DE genes during the night (336) than during the day (233; **SI Table 4**). Large numbers of DE transcripts were unique to each CAM taxon, indicating abundant lineage-specific changes (**Fig. 4A**). As expected, ***Tfasc***, the species with the strongest CAM phenotype and the greatest phylogenetic distance to the C3 reference taxon, exhibited by far the greatest number of unique DE transcripts (1838 during the day and 1881 during the night; **Fig. 4A**).

**Figure 4.**
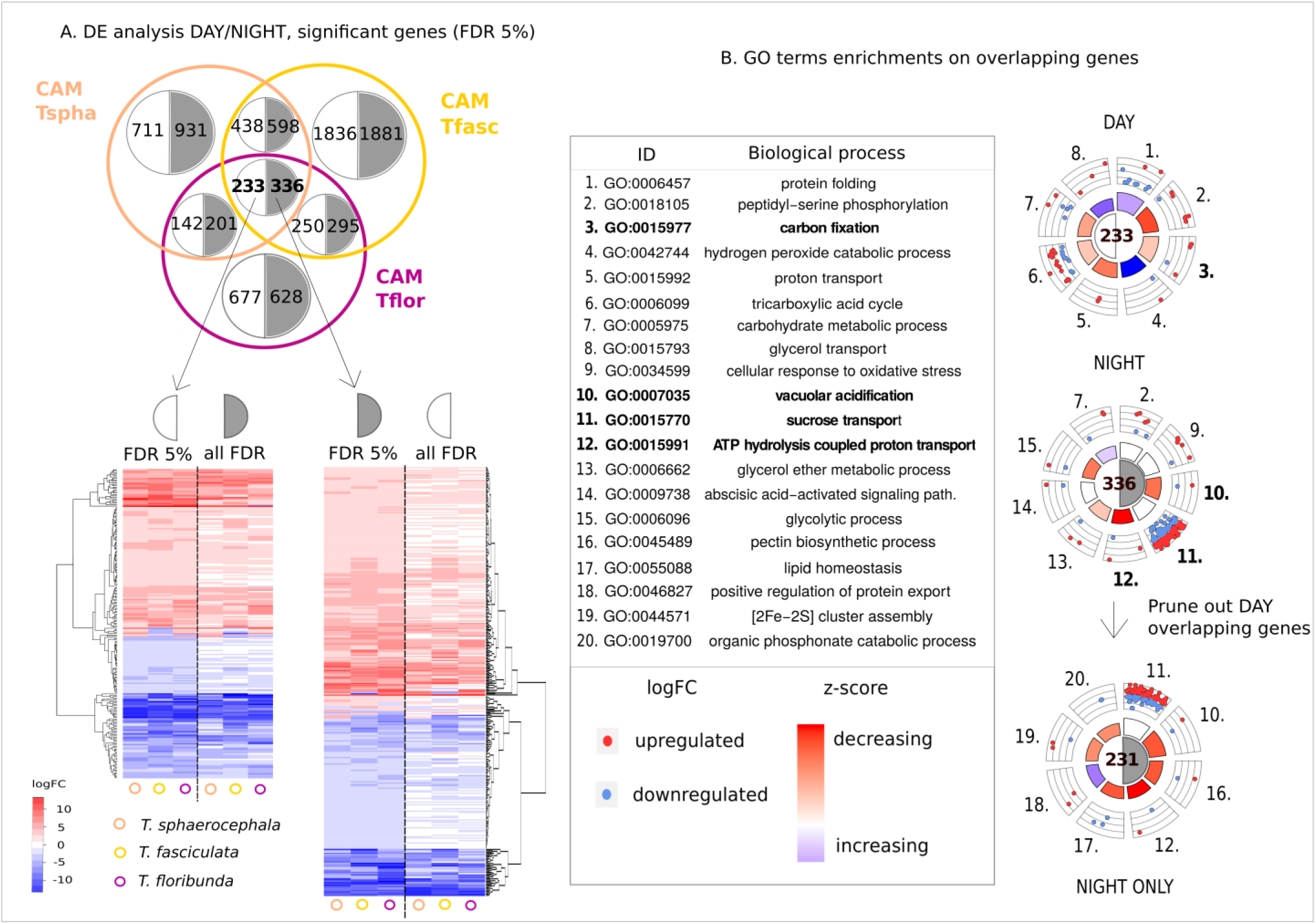
**A**, Transcriptome-wide analysis of differential gene expression (DE) between three CAM Tillandsia species and a C3 reference taxon. Top: Venn chart depicting overlap in DE patterns for each of the three CAM taxa relative to the C3 reference taxon during the day (11AM, numbers in light half-circles) and during the night (1AM, numbers in grey-shaded half-circles). Bottom: Heatmaps illustrating expression changes for the 233 and 336 transcripts with DE in all three CAM species at day and night, respectively, compared to the same set of transcripts at the respective alternative sampling time point (night and day). The similarities in expression patterns of most DE transcripts between day and night are clearly visible. **B**, GO enrichment analysis on the subsets of overlapping significant genes during the day (GO terms n° 1 to n° 8), during the night (n°2, n°7, and n° 9 to n° 15), and a subset of night-specific genes after pruning out the day-overlapping genes. For each subcategory (day, night, and night only) we depict the 8 most significantly enriched GO terms, with emphasis on the photosynthesis-related day-specific (bold, n°3) and night-specific (bold; n°10 to n°12) ones. Rosette plots highlight the relative contribution of up- and downregulated genes to each term and the overall trend (middle circle).

Genes with DE in interspecific C3/CAM comparisons common to all three CAM species were enriched for a range of CAM-related metabolic processes in photosynthetic leaf tissues (Fig 3B). For example, day time points were enriched for genes involved in carbon fixation (GO:0015977), carbohydrate metabolic process (GO:0005975), and the TCA cycle (GO:0006099). In turn, DE genes at night showed enrichment for vacuolar acidification (GO:0007035), sucrose transport (GO:0015770) and glycolysis (GO:0006096), among others. These results provide evidence for large-scale transcriptional reprogramming of CAM-related pathways shared among CAM ***Tillandsia*** species. A more detailed exploration of regulatory changes shaping carbohydrate metabolism based on KEGG maps is given in Supplementary Materials (**SI Text, SI Fig. 6ABC**).

Among the common CAM/C3 DE genes (up-regulated in all CAM compared to the C3 species) there are also several genes involved in abscisic-acid signalling (GO:0009738), including Aco005513 a homolog of ***AtPYL***, an ABA receptor involved in drought response and leaf senescence in ***A. thaliana***^24^, and Aco004854, a further member of the ABF2/ABF3 gene family involved in ABA-mediated response to sugar, drought and osmotic stress^32^. DE genes also included five LEA genes (**SI Table 4**) likely involved in drought protection^33^ and Aco004804, a homolog of the ***A. thaliana*** gene ***YLS7*** identified as a QTL for drought resistance in this species^34^. Together, these results place a strong emphasis on the importance of drought response to the evolution of the CAM-related adaptive syndrome.

### Targeted DE analysis for CAM-related genes from pineapple

Available functional annotation and gene expression information from ***A. comosus***^18,35^ facilitated targeted analysis of known CAM-related genes in ***Tillandsia*** spp. and comparisons to pineapple. Based on published information, we identified 35 homologue clusters of putatively CAM-related genes in the annotated ***A. comosus*** reference genome (**Fig. 5; SI Figs. 7 and 8**). The results revealed diel cycling and upregulation in CAM species of the key post-translational regulator of CAM photosynthesis, phosphoenolpyruvate carboxylase kinase (PPCK, also commonly referred to as PEPC kinase, ***A. comosus*** gene model Aco13938; **Fig. 5**). We also detected significant upregulation of a PEPC gene (Aco010025) in all three CAM ***Tillandsia*** spp., indicating that this is the homolog involved in nocturnal carbon fixation, consistent with its proposed function in pineapple^35^ (**Supplementary information**). At a broader level, targeted DE analysis of known pineapple CAM genes mirrored important patterns recovered from the global functional enrichment analysis (**above; Fig. 4B**). These include upregulation of malate transporters and vacuolar proton pumps (enabling metabolite transport) in ***Tillandsia*** CAM species relative to our C3 reference ***(Taust)*** (**Fig. 5D**), and significant expression changes in many transcripts involved in glycolysis and gluconeogenesis (**SI Figs. 6B** and 7), two important pathways in photosynthetic leaf tissues of plants. Also, starch synthase and other enzymes of starch metabolism were upregulated in CAM plants (**SI Fig. 6C and 8**), consistent with the use of starch as the predominant transitory carbon storage compound in ***Tillandsia*** spp.^22^, as opposed to sucrose in the case of pineapple^18^.

**Figure 5.**
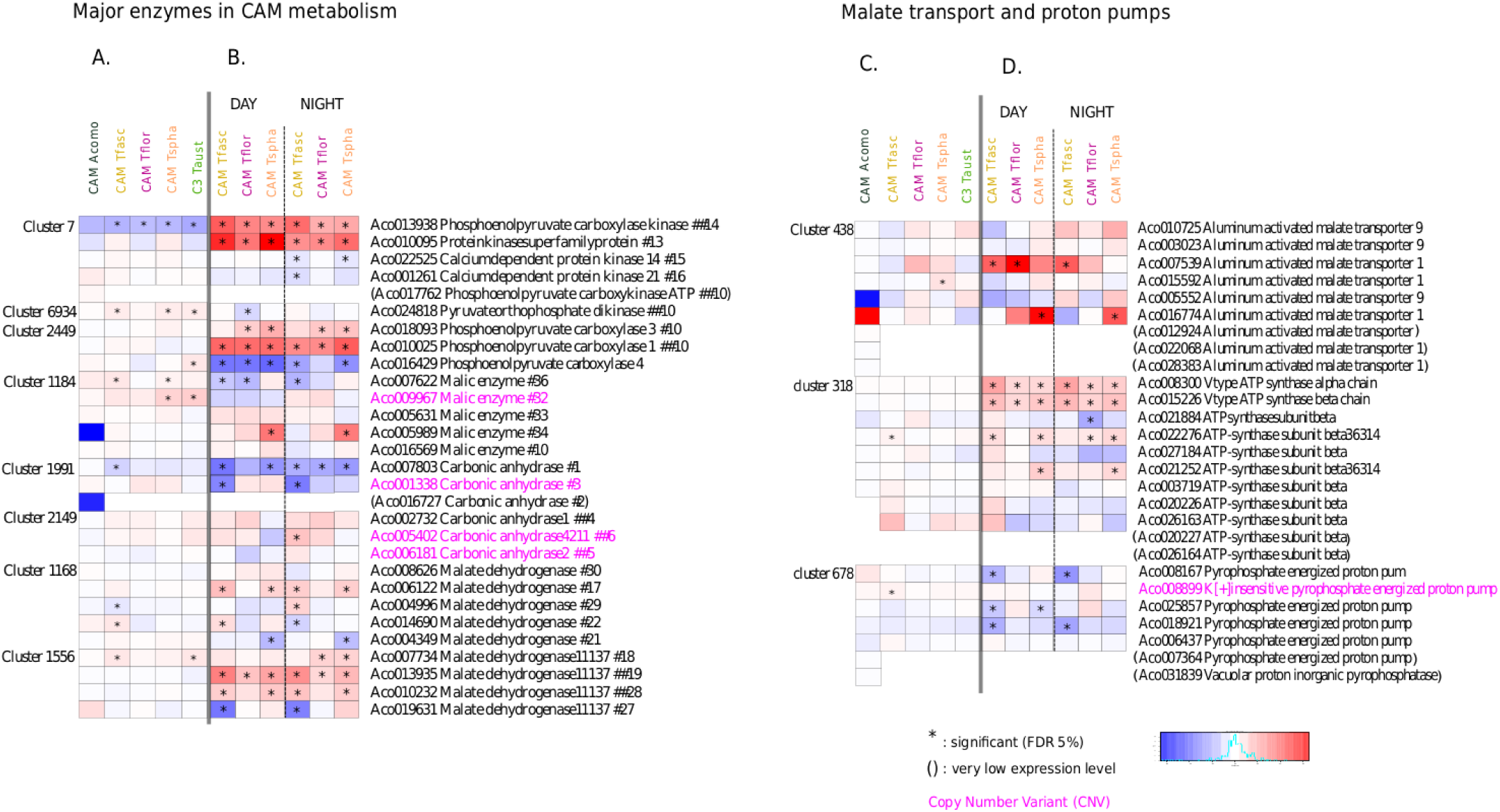
Heatmap of 35 *A. comosus* (pineapple) candidate genes and homologs. Exemplary heatmaps depicting DE patterns for homologue clusters of genes with known or suspected involvement in CAM photosynthesis based on evidence from *A. comosus* (Acomo)^18,33^. Species identities, abbreviations, and color codes are as in **Fig. 2**. **A**, DE for homologue clusters surrounding Acomo core CAM pathway genes for intraspecific day / night tests, including expression information for Acomo for the same time points^18^. **B**, DE for the same homologue clusters in interspecific C3 / CAM tests for the day and night sampling time points. **C**, DE for homologue clusters surrounding genes for malate transport and vacuolar proton pumps for intraspecific day / night tests, including expression information for Acomo for the same time points^18^. **D**, DE for the same homologue clusters in inter-specific C3/CAM tests for the day and night sampling time points.

## Discussion

Our whole-genome phylogenomic data point to extensive associations between CAM and numerous other plant traits linked to xeric adaptation, including leaf succulence, trichomes, leaf shape, and rosette morphology (**Fig. 1**), as previously observed by evolutionary biologists^9–11^ and plant physiologists^16,17,36^. Combining genomic signatures with transcriptomic and targeted metabolite analyses, we revealed the genetic and molecular components underpinning this correlated adaptive trait syndrome in tillandsioid bromeliads.

Together, our data paint a picture of complex interactions of evolutionary changes to gene expression levels, metabolite-based regulation and physiological transitions. We discovered convergent regulatory changes shared among the three CAM species for hundreds of genes (**Fig. 4**; **SI Table 4**), including significant upregulation of malate transporters and vacuolar proton pumps (**Fig. 4D**), commonly associated with increased leaf succulence^16,17,22^, and extensive upregulation of transcripts involved in carbohydrate metabolism and fluxes (**SI Fig. 8**) as expected for CAM plants^37^. Widespread convergence in gene expression patterns accompanying C3/CAM shifts suggests that the most plausible evolutionary path would be the alteration of a shared regulator of these gene sets and that the regulatory networks active in CAM species are largely already in place in C3. Indeed, we found changes in gene expression and/or gene family expansion for several TFs and a sensor involved in ABA signalling, promising candidates for regulators mediating this transition. We also discovered convergent expansion of the XAP5/XCT gene in several CAM lineages (**Fig. 3**, right panel). This gene encodes a highly pleiotropic regulator known to be involved in a variety of processes including light-dependent gene expression and developmental processes in ***A. thaliana***^28–30^. CAM genes are regulated in a circadian-clock dependent manner in the bromeliad ***A. comosus***^35^ and duplication of XCT could have helped to mediate light-based reprogramming of transcription in CAM species. Gene duplication was also detected for a galactinol synthase gene which may play a role in drought resistance^27^, underlining this to be a common theme among different evolutionary mechanisms captured by our dataset.

Besides pleiotropic TFs, we also found indications for alterations to genes involved in regulation at the metabolic level. In particular, branch-specific tests for selection highlighted the enolase Aco020962, a homolog of AtENO2/LOS2. This gene is of particular interest to correlated evolution involving CAM, drought response, and carbon fluxes, since it provides a direct link between glycolysis/gluconeogenesis and the CAM pathway via PEP, catalysed by the enolase-coding isoform, and ABA-dependent response to abiotic stress mediated by the regulatory protein MBP-1 transcribed from an alternative start codon at the same locus^38^. Notably, this gene exhibited significant temporal and interspecific C3/CAM expression changes in *T*. ***fasciculata (Tfasc)***, the strongest CAM species in our transcriptome study. CNV data indicate this gene was present in multiple copies in ***Tillandsia*** genomes, and it is thus possible that increased expression in the strong CAM plant ***Tfasc*** and a high estimate of non-synonymous substitutions was due to recent gene duplication, hypotheses that remain to be tested in the future.

A persistent pattern recovered by both global and targeted gene expression profiling is the general absence of a clear temporal signature of diel cycling of CAM genes that distinguishes CAM- and C3-like species (**Fig. 4**). Instead, we found sweeping regulatory changes to key CAM genes and many other pathways relevant for the CAM phenotype, often evident during both day and night (**Figs. 4 and 5**). Consequently, our data suggest that rather than altering the timing of expression, relevant genes already follow CAM-like expression patterns in the C3 species but their expression is amplified in CAM species (**Fig. 4**). These observations provide mechanistic insights into the evolutionary mechanisms supporting the C3/CAM continuum^16,36^ and suggest that genes already following a CAM-like expression pattern are preferentially co-opted into the CAM pathway, as suggested by Bräutigam et al (2017)^39^. This may explain the predominance of convergent co-option of the same homologs among our three CAM species surveyed (**Fig. 5**) and mirrors observations made in C4 grasses, where recurrent co-option of specific C4 homologs in independent origins of C4 photosynthesis was driven by expression levels of the respective gene copies in photosynthetic leaves of the C3 ancestor^40^.

In summary, our combined whole-genome, transcriptome, and metabolic data point to a central role of highly pleiotropic regulators and drought response-related pathways to the repeated origin of CAM photosynthesis in tillandsioid bromeliads, involving both “top down” and “bottom up” changes. We thus hypothesize that transitions to CAM photosynthesis involved the re-programming of drought-related pathways. We think that the striking correlated trait shifts seen in this textbook adaptive radiation are due to pleiotropic regulators, rather than genomic clustering of adaptive mutations. We expect this view to be corroborated or challenged by genomic data to emerge in years to come.

## Materials and Methods

### Species sampling

Specimens were sampled from the living collections of the Universities of Vienna (Austria) and Heidelberg (Germany) and in the Botanical Gardens of Rio de Janeiro (Brazil), Lyon (France), Porrentruy, and Geneva (both Switzerland). Information on morphological characters, habitats, and natural distributions of the studied accessions is provided in **Fig. 1** and **SI Table 1**.

### DNA extraction and whole genome sequencing (WGS)

For each sample, 30mg of leaf material was dried in silica gel before DNA extraction with a QIAGEN DNeasy® Plant Mini Kit, following supplier’s instructions. After shearing of 1μg of DNA with either a Covaris® or Bioruptor® instrument, sequencing libraries were prepared with the Illumina TruSeq® DNA PCR-Free Library Prep Kit. Libraries were sequenced paired-end 2×150bp or 2×125bp in seven different lanes of an Illumina HiSeq3000 sequencer.

### WGS data processing and variant calling

Reads were trimmed with condetri v2.2^41^ using 20 as high-quality threshold parameter. The *Tillandsia adpressiflora* sample with the highest number of reads (**SI Table 1**) was used to build a pseudo-reference genome following an iterative mapping strategy described in de La Harpe et al. (2018)^42^, using the annotated *A. comosus* reference genome^35^ as a starting point. The 28 samples and *A. comosus* were mapped to the pseudo-reference using Bowtie2 v2.2.5^43^ with the very-sensitive-local option. SNPs were called using GATK UnifiedGenotyper using the EMIT_ALL_SITES and –glm SNP options, after realignment around indels and base recalibration with GATK v3.3^44^. Positions were filtered with vcftools v0.1.13^45^ before subsequent analyses retaining only positions with quality >20, read depth >3 and a maximum of 50% of missing data. Read number, coverage and mapping statistics were calculated using bedtools v2.24.0^46^ and vcftools v0.1.13^45^.

### Phylogenomic analysis

In order to reconstruct evolutionary relationships among the target species, we used ASTRAL v5.6.1^47^, a summary method based on the multispecies coalescent, to infer a species tree (**Fig. 1; SI. Fig. 1**), complemented by maximum likelihood analysis in RAxML v8.228^48^. A phylogenetic network was constructed using the neighbour-net method implemented within SplitsTree^20^ using an identity-by-descent distance matrix obtained with PLINK^49^.

### Carbon isotope phenotyping

We assessed the carbon isotope ratio (^13^C/^12^C) for all species used in the present study (**Fig. 1**) using 1 gram of silica-dried material per sample. The measurements were carried out at the Institute of Earth Surface Dynamics (Faculty of Geosciences and Environment, University of Lausanne, Switzerland) following Spangenberg et al. (2006)^50^. This approach makes use of flash combustion on an elemental analyser connected to a ThermoQuest/Finnigan Delta S isotope ratio mass spectrometer via a ConFlo III split interface. The carbon isotope ratios values (δ) correspond to the per mille (‰) deviation relative to the Vienna-Pee Dee belemnite standard (V-PDB).

### Targeted metabolite analysis

Dry silica samples were collected in the Bromeliaceae research greenhouses at Department of Botany and Biodiversity Research, University of Vienna, in July 2017 at distinct time points, namely at 1AM and 11AM, following the same day/night sampling scheme also used for gene expression profiling based on RNA-seq (above). We sampled green tissue from mature leaves, where possible from the exact same specimen also used for whole genome sequencing (WGS) and RNA-seq. In four cases for which the exactly same species were not available in the greenhouse, we sampled species known to be very closely related to the actual target taxa. These are denoted with an asterisk (*) in **Fig. 1**. All samples were processed at the Vienna Metabolomics Center (VIME, Department of Molecular Systems Biology, University of Vienna). All analysis steps including plant metabolite extraction, sample derivatization, and GC-TOF-MS (gas chromatography coupled with time-of-flight mass spectrometry) were carried out as previously described^51^. Data analysis was performed using ChromaTof (Leco) software. Briefly, representative chromatograms of different samples were used to generate a reference peak list, and all other data files were processed against this reference list. Deconvoluted mass spectra were matched against an in-house mass spectral library. Peak annotations and peak integrations were checked manually before exporting peak areas for relative quantification. Metabolite amounts are given in arbitrary units corresponding to the peak areas of the chromatograms. Data were analyzed using Partial Least Squares (PLS) Discriminant Analysis (DA) using the the COVAIN toolbox for metabolomics data mining^52^.

### Positive selection on coding sequences

We tested for positive selection acting on the WGS data using the non-synonymous to synonymous substitution rate ratio (ω = dN/dS) tests using codeml in PAML version 4.8a^53^. We filtered out gene alignments with less than 300bp, more than 50% missing data, and multiple stop codons. Additionally, individual sequences with more than 90% missing data were removed from the codon alignment. We inferred gene phylogenies for each codon alignment using PhyML version 3.3.20170119^54^ with HKY and GTR substitution models and GAMMA shape parameter. We computed a Likelihood Ratio Test (LRT) to select the best reconstruction for codeml analyses. We ran the cladeC model containing five estimated parameters denoted: p0, p1, p2 (p2= 1 – p0 – p1) for the proportion of sites in a site-class, and dN/dS parameters ω0, ω1 fixed to 1, ω2 background (C3 plants), and ω2 foreground (CAM plants). This model tested whether selection has occurred in all branches leading to species with CAM metabolism (H1). The null model corresponded to the M2a-rel^55^ with four parameters, where ω2 background and foreground are fixed to be equal. Each model was run three times to overcome convergence issues, and the best likelihood run was used for model comparison and ω estimates. Model fit was compared using the corrected Akaike Information Criteria (AICc) with a significance threshold delta-AIC of 10 between M2a-rel and H1. For genes preferring H1, we used the standard errors (SE) of ω estimates to determine the deviation from neutrality (ω=1 is nearly neutral, therefore genes were discarded when ω ± SE included 1), and considered only genes with signatures of positive selection in any CAM lineage (ω2 in CAM > 1). Genes with dN/dS estimates close to the optimization bound in the three replicates (i.e. parameter estimates close to 999) and SE larger than 10 were discarded.

### Inference of gene family evolution

Copy number variants were detected in 15 high-coverage individuals (Fig. 3) based on relative read-depth differences in exons using CNVkit^56^. Illumina reads were aligned to the *Tillandsia* pseudo-reference as above and filtered using SAMtools v.0.1.19^57^ to remove duplicates (rmdup) and ambiguously mapping reads with mapping quality less than 10 (-q 10). To avoid conflating lack of mappability with gene loss, we only retained exons with a mean read coverage of at least five in five individuals for further analysis. We first called copy number (CN) status in *Al. trepida* by profiling against *A. comosus* as a reference. For CNV detection within the genus *Tillandsia*, we then used *A. trepida* as the reference sample since it better reflects the observed mapping biases against the pseudo-reference genome with the remaining *Tillandsia* samples (lack of coverage outside coding regions, systematic variation in coverage along the genome). CNVkit was run with an average antitarget size of 5000 and an accessibility mask of 2 kb generated based on the *A. comosus* hardmasked reference sequence available from CoGe (https://genomevolution.org/CoGe/GenomeInfo.pl?gid=25734). Gene gains and losses were called requiring consistent signal across at least four tiles (gainloss -m 4). Log2 ratios per gene were then filtered according to coverage-dependent, empirically determined thresholds for lower and upper bounds (Supplementary Text). Log2 ratios above and below the respective thresholds were then translated into numbers of alleles by exponentiation and multiplication with the inferred allele number in *A. trepida*, divided by two and rounded to the next integer in order to calculate CNs per gene. We consider CNs to be an approximate and relative measure of CN variation with the genus *Tillandsia* rather than absolute estimates.

Homolog clusters in the pineapple were identified using MCL clustering of the *A. comosus* predicted proteins with MCL-edge^58^, based on an all-vs-all BLASTp search with an e-value cutoff of 1e^-5^ and an inflation value of 3. Inferred CNs for each gene in a cluster were summed to obtain family-level gene pseudocounts across the high-coverage dataset for analysis of gene family evolution with CAFÉ v4.1^25^. We pruned species not included in the CNV analysis from the RAxML tree and used the Penalized Likelihood method with cross validation implemented in the program r8s^59^ to obtain an ultrametric tree for analysis with CAFÉ. All CAFÉ models were run with species-level error models inferred using the error model estimation script supplied with CAFÉ. Error estimates by species ranged between 0 in *A. trepida, T. sphaerocephala* and *T. hitchcockiana* to a maximum of 0.03 in *T. juncea* and *T. leiboldiana*. We estimated separate rates of gain (λ) and loss (μ) in either a global model with a single set of parameters across the entire tree or a two-rate model with different sets of rate parameters for species in the subgenus *Tillandsia* and the rest of the tree (Fig. 3). The two-rate model fit significantly better than the global model (delta-AIC 1727) and was the basis for inference of gene gains and losses along branches and ancestral copy numbers.

To test for associations between CN changes and evolution of CAM photosynthesis and correlated traits we ran a permutation ANOVA^60^. Branch-wise turnover rates were calculated from ancestral CN estimates for each family and branches were labelled according to photosynthetic strategy. Empirical P-values were determined from 1000 permutations of branch label swapping and corrected for multiple testing to control the False Discovery Rate^61^.

### RNA sequencing

All four species were kept under identical greenhouse conditions for at least 10 days prior to sampling (°C: min=16.6, max=39.0, mean=25.7 and %rF min= 30.8, max=95.9, mean=67.5). Three different specimens were used for each species to serve as biological replicates, except for *Tflor* for which only two replicates were available. For each sample and time point, up to 30mg fresh leaf tissues were stabilized in RNAlater® immediately after sampling and kept at −20°C. Total RNA was carefully extracted under a sterilized fume hood with the QIAGEN RNeasy® Mini Kit following the supplier’s protocol. RNA libraries were prepared with the Illumina TruSeq® Stranded mRNA Library Prep Kit before sequencing pair-end 2×150bp on an Illumina HiSeq3000 sequencer.

Sequence quality was validated with FastQC v0.11.2 (https://www.bioinformatics.babraham.ac.uk/projects/fastqc/README.txt), and reads were trimmed with condetri v2.2^41^ using 20 as high-quality threshold parameter. Reads were mapped to a *Tillandsia adpressiflora* pseudo-reference genome (described below) with TopHat v2.1.0^62,63^ using *--b2-very-sensitive* mapping parameters. Only uniquely mapped reads were kept for further analyses.

### Differential expression analysis of RNA-seq data

The number of reads mapping to reference genes was quantified with HTSeq-count in default mode^64^, producing the gene count data (27,024 genes in total) for the DE analyses. We filtered the database by removing genes with 1 and 0 counts, resulting in a database of 23,737 genes. We used edgeR v.3.12.1^65^ for DE analysis. After filtering out lowly expressed genes (cpm < 2) we constructed a General Linearized Model (GLM) with species and time point as factors to implement the following tests: (1) “Intraspecific day/night” to detect genes with DE between day (11AM) and night (1AM) time points within each species, and (2) “Interspecific C3/CAM” to detect DE genes between CAM and C3 species at the day and night time points respectively. Significance was tested using likelihood ratio tests and P-values were corrected for multiple testing using the FDR. To avoid potential complications arising from copy number variation, we removed genes with evidence for copy number variants (CNVs; above) before further analysis unless otherwise noted. Gene Ontology (GO) terms were extracted for 20'165 genes (74.61% of the annotated genes) with Blast2GO^66^ in April 2017. GO enrichment analyses were performed with the R package topGO v.2.22.0^67^ using Fisher’s exact test and the weight01 algorithm. GOplot^68^ was used to graphically represent the significantly enriched GO terms and visualize enrichment for the ‘biological processes’ domain by calculating the z-score as z = (up-down)/ Vcount).

## Supporting information

## Acknowledgements

We thank Michael Kessler and other members and collaborators of Swiss SNSF Sinergia project CRSII3_147630 for helpful discussions; the SNSF for funding; staff of all living collections and service facilities used; and Jim Leebens-Mack and Karolina Heyduk for commenting on earlier versions of the paper.

## Author contributions

CL, MDLH, MP, JH, and NS conceived the study, CL and NS provided funding, MDLH, MP, MHJB, AG, and WT collected data, MDLH, MP, JH, MLSS, PC, and WW analyzed data, and MDHL, JH, and CL wrote the paper with input and revisions from all coauthors.

## Competing interest statement

The authors declare no competing interests

## Data availability

The datasets generated during and/or analysed during the current study are available from the corresponding author on reasonable request. Sequence reads will be deposited in NCBI-SRA (accession ID XXXX).

