## Supplementary material for "Genomic footprints of repeated evolution of CAM photosynthesis in tillandsioid bromeliads"

[1] University of Vienna, Faculty of Life Sciences, Department of Botany and Biodiversity Research, Division of Systematic and Evolutionary Botany, Rennweg 14, A-1030 Vienna, Austria, [2] University of Fribourg, Department of Biology, Unit of Ecology & Evolution, Chemin du Musée 10, CH-1700 Fribourg, Switzerland, [3] University of Lausanne, Faculty of Biology and Medicine, Department of Ecology and Evolution, Biophore, 1015 Lausanne, Switzerland, [4] University of Vienna, Faculty of Life Sciences, Department of Ecogenomics and Systems Biology, Division of Molecular Systems Biology, Althanstraße 14 (UZA I), 1090 Vienna, Austria, [5] University of Vienna, Vienna Metabolomics Center (VIME), Althanstraße 14 (UZA I), 1090 Vienna, Austria, [6] Michigan State University, College of Agriculture & Natural Sciences, Department of Horticulture, 1066 Bogue Street, East Lansing, MI 48824, U.S.A, [7] University of Illinois at Urbana-Champaign, School of Integrative Biology, Department of Plant Biology, 265 Morrill Hall, MC-116, 505 South Goodwin Avenue, Urbana, IL 61801, U.S.A.

#### Supplementary Methods

##### 1) Carbon isotope phenotyping

Whole tissue carbon isotope ratios ($^{13}C/^{12}C$) can be used to characterize typical C3 and CAM plants[1], especially in combination with distinct morphological and anatomical features as are present in bromeliads (**Fig. 1**). Carbon isotope ratios recovered for the studied species indicate a continuum of values ranging from typical C3 to fairly strong CAM (**Fig. 1**), following commonly used thresholds[2,3]. Many species in our sample set displayed typical C3 carbon isotope ($\delta^{13}C$) phenotypes far beyond -20‰ and in fact reaching as far into the C3 extreme as -30‰ (labelled green in **Fig. 1**). *Tillandsia australis (Taust)*, our C3 reference taxon also used for transcriptome-wide expression profiling (below), exhibited $\delta^{13}C$ values of -26 to -29‰, clearly beyond the -20‰ threshold commonly used to classify C3 plants[1]. This species also exhibited all other phenotypic features expected for C3 bromeliads, including tank-forming rosettes, no succulence, and absence of dense trichome cover. On the other end of the C3/CAM continuum, *T. ionantha* exhibited a $\delta^{13}C$ value of only -13.9‰, indicating it represents a so-called ′strong′ CAM species (labelled in yellow in **Fig. 1**). The three CAM taxa sampled for expression profiling in our study (below) exhibited a broad range of CAM-like $\delta^{13}C$ values, from weak CAM in *T. floribunda* (*Tflor*, $\delta^{13}C$ = -18.6 to -20.3‰) to relatively strong CAM in *T. sphaerocephala* (*Tspha*, $\delta^{13}C$ = 15.2 to -16.2‰) and *T.*

*fasciculata* (*Tfasc*, $\delta^{13}C$ = -14.5 to -18.3); species exhibiting pronounced, strong CAM typically exhibit $\delta^{13}C$ values that are less negative than -20‰[1].

Although measuring night-time acidity under drought stress is preferable for distinguishing true C3 species from inducible, facultative CAM species[1,4], we opted for carbon isotope ratios based on clear restrictions presented by our study system. First, our sampling of highly divergent phenotypic forms made it challenging to derive comparable 'common garden' drought conditions across all species. Second, our work relied on precious living collections in botanical gardens and experimentation thus required careful consideration to avoid the loss of individual accessions. While we cannot exclude that some species classified as C3-like here include facultative CAM plants, we use isotopic ratios as a proxy to partition species according to the extremes of the distribution of this continuous phenotypic trait for our evolutionary analyses. This is a conservative and pragmatic strategy, since phenotyping error would likely diminish the signal-to-noise ratio.

#### 2) Phylogenetic analyses

We used two complementary methods for species tree estimation: ASTRAL, a coalescent-based summary method[5] (**Fig. 1**) and RaxML which infers maximum likelihood (ML) based phylogenetic trees (**SI Fig. 1**). The ML tree was inferred using the program RAxML v8.228[6] with a GTRGAMMA model and 100 bootstrap replicates to determine branch support. For detailed settings used for estimation of the ASTRAL tree please refer to Materials and Methods in the main text.

Both trees are identical, except the position of *Tillandsia disticha* which in the ASTRAL tree is placed sister to subgenus *Tillandsia.* In the RAxML tree this species is inferred as basal to all main *Tillandsia* divisions.

#### 3) CNV analyses: Detailed implementation of the CNVkit analysis

Relative copy numbers were estimated in a two-step approach: i) estimation of base copy number (CN) state in *Alcantarea trepida* through comparison with *Ananas comosus.* ii) estimation of CN in *Tillandsia* samples relative to *A. trepida,* scaling of Tillandsia CN with *A. trepida* base CN. For both analyses we applied a stringent coverage-based filtering to exclude unmappable regions since we cannot distinguish them from lost genes. Coverage cutoffs were set as follows: filtered_exon_set_1 (Aco-Atre analysis) – retain only exons with mean coverage of at least five, filtered_exon_set2 (Atre-Tillandsia analysis) – retain only exons with mean coverage of at least five in five or more species. This resulted in a total of 19,298 discoverable genes in filtered_exon_set2 (roughly 2/3rds of the genome).

In order to derive meaningful log2 thresholds for CNV calling, we used deeply conserved single-copy orthologs in the pineapple to estimate variation in coverage. To this end, we used BUSCO v3[7] to obtain the set of single-copy orthologs using the *A. comosus* predicted proteome and the "embryophyta_odb9" database. To derive per-sample single copy thresholds we calculated the weighted average log2 ratios across exon bins for each of the BUSCO genes. The resulting distributions are shown below.

| **Species** | **1st Qu.** | **Median** | **Mean** | **3rd. Qu.** | **2.50%** | **97.50%** |
|---|---|---|---|---|---|---|
| *Tillandsia leiboldiana* | -0.1389 | -0.02263 | -0.01518 | 0.09641 | -0.4906562 | 0.5907426 |
| *Tillandsia australis* | -0.08921 | -0.003557 | -0.004812 | 0.07833 | -0.4050659 | 0.4650078 |
| *Tillandsia propagulifera* | -0.08921 | -0.003557 | -0.004812 | 0.07833 | -0.3229344 | 0.3847263 |
| *Tillandsia floribunda* | -0.09287 | -0.003627 | -0.00184 | 0.07432 | -0.3525346 | 0.3999353 |
| *Tillandsia latifolia* ssp. *latifolia* | -0.1014 | -0.01013 | -0.006266 | 0.08504 | -0.4294942 | 0.6181706 |
| *Tillandsia trauneri* | -0.1078 | -0.008357 | 0.005459 | 0.0832 | -0.3475898 | 0.6060711 |
| *Tillandsia hitchcockiana* | -0.1047 | -0.01453 | -0.006099 | 0.07408 | -0.3488612 | 0.5124046 |
| *Tillandsia sphaerocephala* | -0.09096 | -0.009717 | 0.001125 | 0.07442 | -0.3531745 | 0.4825815 |
| *Tillandsia adpressiflora* | -0.08112 | -0.007189 | -0.01488 | 0.06597 | -0.3108748 | 0.4262556 |
| *Tillandsia somnians* | -0.08239 | -0.005033 | -0.005541 | 0.07851 | -0.3314174 | 0.550001 |
| *Tillandsia stenoura* | -0.09341 | -0.004734 | 0.02142 | 0.09441 | -0.3748035 | 0.6693527 |
| *Tillandsia complanata* | -0.08054 | -0.0008737 | 0.02569 | 0.08393 | -0.2753471 | 0.568465 |
| *Tillandsia fasciculata* | -0.08956 | -0.01163 | 0.007404 | 0.07773 | -0.336556 | 0.5267081 |
| *Tillandsia juncea* | -0.09474 | -0.007876 | -0.004926 | 0.06462 | -0.3213492 | 0.4067321 |
| *Tillandsia stricta* | -0.1053 | -0.005714 | -0.0399 | 0.0946 | -0.4515171 | 0.4965152 |

| | | | | | | |
|---|---|---|---|---|---|---|
| *Vriesea itatiaiae* | -0.09843 | -0.01375 | -0.01111 | 0.06589 | -0.3339326 | 0.3725004 |

Based on these results, we settled to set cutoffs of log2(allele_count-0.5/alleles) for copy number decrease and log2(allele_count+1/alleles) for copy number increase which corresponds to log2(1.5/2) and log2(3/2) for a single copy locus with two alleles. This encompasses the empirically observed range of variation and is dynamically adjusted to accommodate increasing variation we expect to be associated with CN > 1 in the reference sequence *A. trepida*.

#### CAFÉ error model estimation and model selection

In order to account for inaccuracies in the CN estimates, as well as differences in accuracy between species, e.g. due to variation in coverage, we estimated an error model to be applied to each species via a built-in estimator supplied with CAFÉ. This resulted in the following best-fit error models:

| **Sample Name** | **Error rate** |
|---|---|
| *Tillandsia fasciculata* | 0.03515625 |
| *Tillandsia trauneri* | 0.0309375 |
| *Tillandsia propagulifera* | 0.00140625 |
| *Tillandsia juncea* | 0.03515625 |
| *Tillandsia latifolia* ssp. *latifolia* | 0.01125 |
| *Tillandsia australis* | 0.00703125 |
| *Tillandsia hitchcockiana* | 4.34E-19 |
| *Tillandsia floribunda* | 0.00703125 |
| *Tillandsia leiboldiana* | 0.06328125 |
| *Tillandsia complanata* | 0.01125 |
| *Tillandsia somnians* | 0.00703125 |
| *Tillandsia adpressiflora* | 0.00140625 |
| *Tillandsia stenoura* | 0.00984375 |

We first ran CAFÉ using a single global rate model. Based on the observation of an apparent increase in the rates of duplication and losses in the subgenus *Tillandsia,* we also tested a two-rate model allowing for separate rates of evolution in this subgenus. Each model was run three times to check convergence of ML estimates and significance was determined using the Akaike Information Criterion (AIC) as below:

| ***Global model*** | $\lambda$ | $\mu$ | **Score** |
|---|---|---|---|
| **Run1** | 0.00101061203283 | 0.00027763568401 | 61203.9 |
| **Run2** | 0.00101060963310 | 0.00027763236492 | 61203.9 |
| **Run3** | 0.00101060978654 | 0.00027763425550 | 61203.9 |
| ***Two-rate model*** | $\lambda$ | $\mu$ | **Score** |
| **Run1** | 0.00079544338562<br>0.00284107269136 | 0.00023883871686<br>0.00086474231103 | 60340.4 |
| **Run2** | 0.00079544582304<br>0.00284121037836 | 0.00023883335117<br>0.00086475802060 | 60340.4 |
| **Run3** | 0.00079543765336<br>0.00284102919941 | 0.00023884156640<br>0.00086464833310 | 60340.4 |

Model testing using AIC with the best run of the one- and two rate models:

| | **lnL** | **#params** | **AIC** | **$\Delta$AIC** |
|---|---|---|---|---|
| **One rate** | -61203.9 | 2 | 122411 | 1727 |
| **Two rate** | -60340.4 | 4 | 120684 | |

AIC = 2k - 2ln(L) where k = number of parameters.

#### 4) RNA sequencing

Between 4,589,500 and 14,227,598 reads from RNA-seq were successfully mapped to the reference genome, corresponding to 56.1% and 57.4% of reads, respectively. Day/night expression information available for *A. comosus*[8,9] allowed us to examine similarities and differences among CAM-related diel expression patterns between *Tillandsia* spp. and the CAM plant pineapple at equivalent time points (**Fig. 4**). Perhaps most obviously, our comparisons indicated diel cycling of the key post-translational regulator of CAM photosynthesis, phosphoenolpyruvate carboxylase kinase (PPCK, also commonly referred to as PEPC kinase, *A. comosus* gene model Aco13938; **Fig. 4**). This important CAM-related gene was significantly upregulated at night in all three CAM *Tillandsia* species studied

(*Tspha, Tflor, Tfasc*), following the pattern expected from the CAM plant *A. comosus (Acomo)* (**Fig. 4A**). Upregulation of PPCK at night was also significant for our C3 *Tillandsia* reference taxon (*Taust*). This is suggestive of pre-adaptation of C3 tillandsioids for the evolution of CAM and mirrors results obtained on C3 *Flaveria* spp., suggesting that PPCK night-time expression may be a conserved pattern among C3 species[10,11]. PPCK was significantly upregulated in interspecific C3/CAM comparisons involving all three CAM *Tillandsia* taxa studied (*Tspha, Tflor, Tfasc*) relative to our C3 reference (*Taust*) (**Fig. 4B**), consistent with the direct involvement of this important protein in CAM photosynthesis in *Tillandsia* spp. We also detected significant upregulation of a PEPC gene (Aco010025) in all three CAM *Tillandsia* spp., indicating that this is the homolog involved in nocturnal carbon fixation, consistent with its proposed function in pineapple[8].

**Suplementary Material**

**SI_Figure_1:** RAxML tree
**SI_Figure_2:** SplitsTree network
**SI_Figure_3:** MDS plot of RNAseq
**SI_Figure_4:** Day/night RNAseq comparisons
**SI_Figure_5:** GO term enrichments
**SI_Figure_6A:** KEGG carbon fixation (ko00710)
**SI_Figure_6B:** KEGG Starch/sucrose (ko00500)
**SI_Figure_6C:** KEGG Glycolysis/gluconeogenesis (ko00010)
**SI_Figure_7:** Extended target gene list intraspecific test with Pineapple expression data
**SI_Figure_8:** Extended target gene list interspecific C3/CAM with Pineapple expression data

**SI Tables**

**SI_Table_1:** Data table (DNA-seq and RNA-seq samples table with morphological characters)
**SI_Table_2:** Table of the metabolite analysis (amino acids, carbohydrates and organic acids)
**SI_Table_3:** Positive Selection Gene list
**SI_Table_4: EXCEL** Table of the DE transcripts (233 during the day and 336 during the night) for the interspecific C3/CAM comparisons.
**SI_Table_5: EXCEL** data table (full-genes list with CNV, DE analysis and positive selection test outputs)

**Supplementary Material for the Tillandsia MS NatPlants**

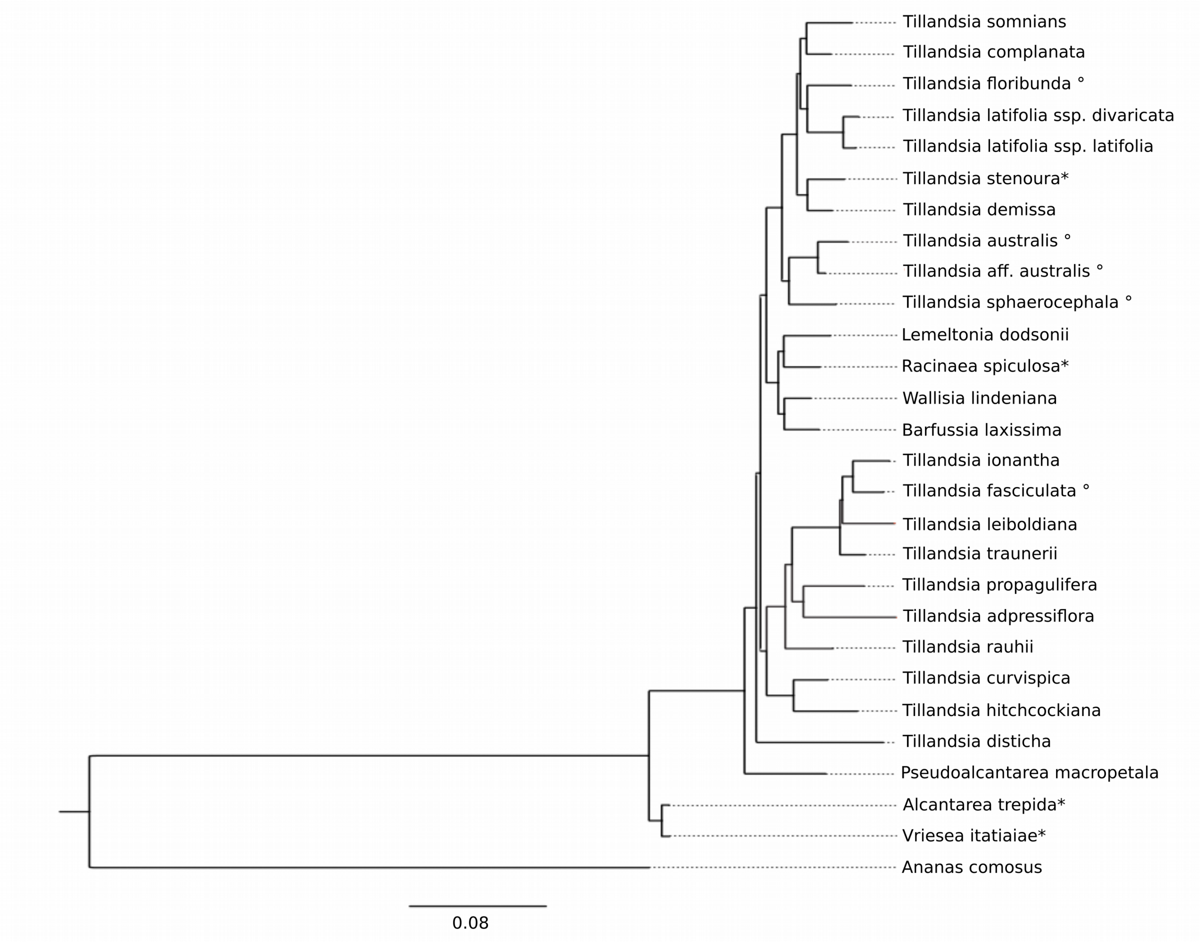


**SI_Figure_1: RAxML** tree : Maximum likelihood (RAxML) phylogenetic tree of 28 whole-genome sequenced tillandsioid bromeliad taxa (species of Tillandsia and related genera), including *Alcantarea trepida* and Vriesea itatiaiae as outgroups for phylogenetic analysis, and Ananas comosus, the species used to anchor most analyses presented in this study.

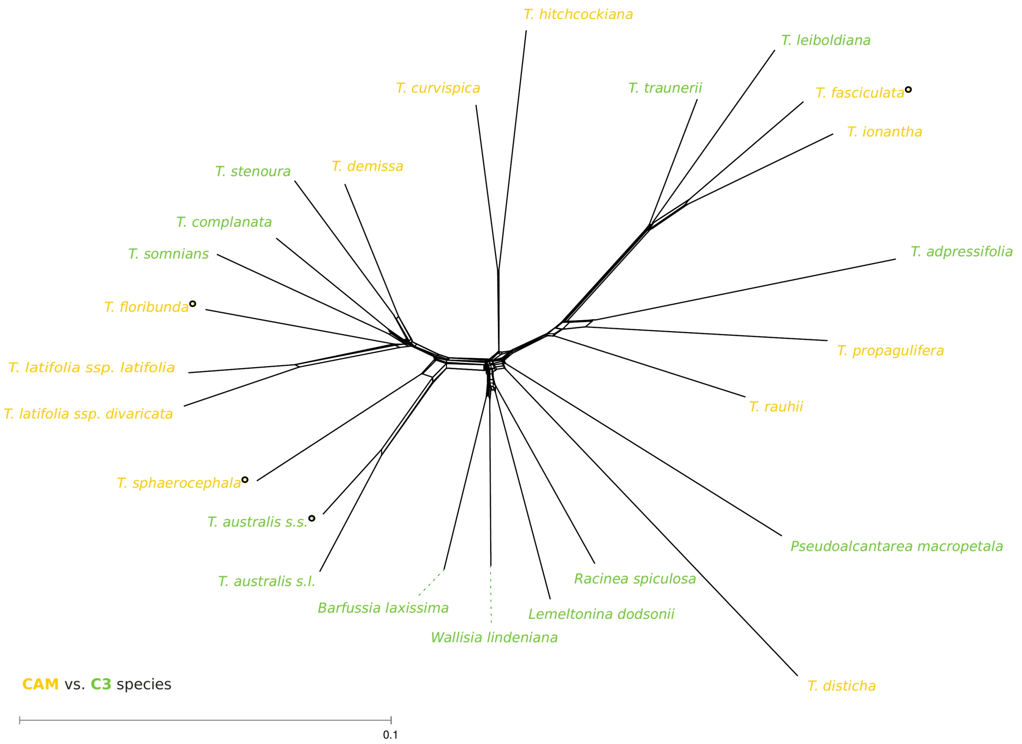


**SI_Figure_2:** SplitsTree network

SplitsTree network based on whole genome sequencing (WGS) of all sampled taxa, with C3 and CAM taxa labelled by different colors.

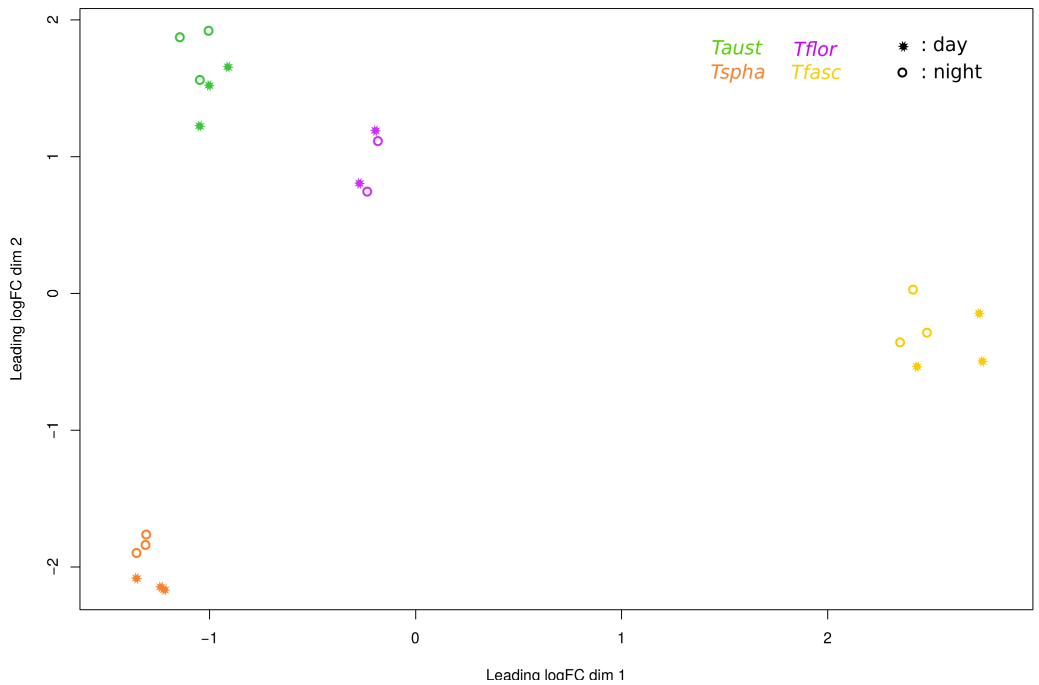


**SI_Figure_3:** MDS plot of RNAseq

**Top axes from Multi Dimensional Scaling (MDS) analysis of differential gene expression (DE) data.**

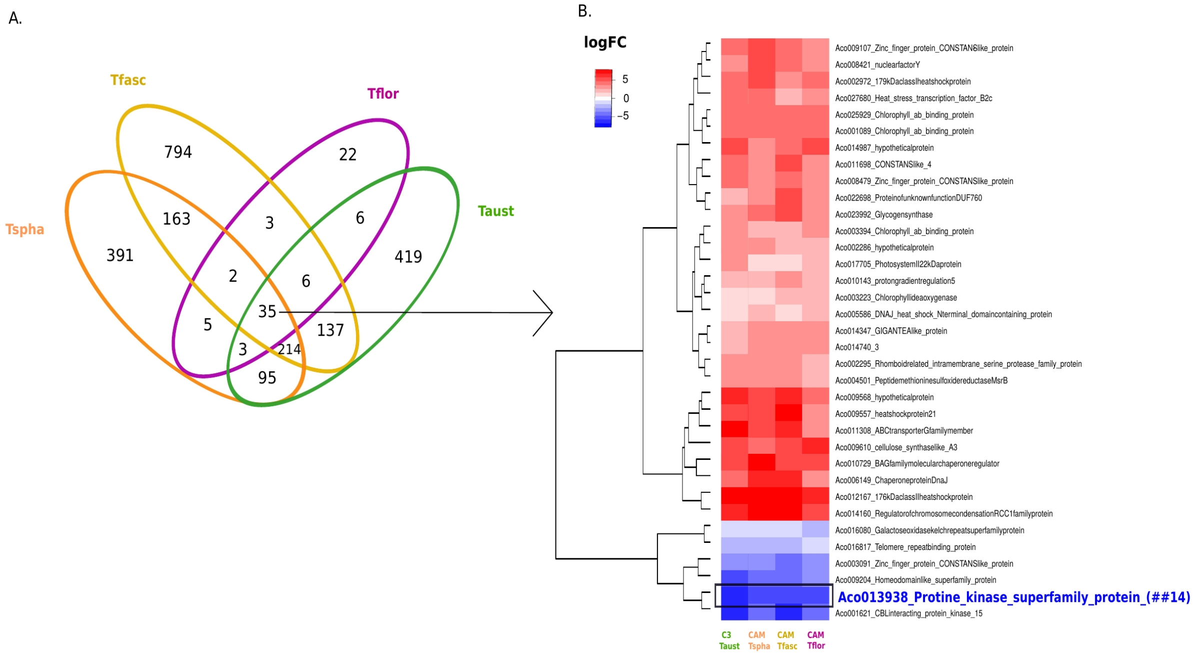


**SI_Figure_4:** Day/night RNAseq comparisons

Results from transcriptome-wide analysis of differential gene expression (DE) between two sampling time points, night (1AM) and day (11AM), referred to as “intraspecific day/night” test throughout the text. Shown are results at FDR<0.05 for logFC values >1 or ←1 (i.e. pruning away logFC values close to zero). A. Venn chart depicting similarities and differences in temporal DE patterns between the studied species. B. Clustering heatmap for transcripts shared by all taxa (35 genes), corresponding to the central overlap field in the Venn chart. Red and blue colors in the heatmap indicate up- and down-regulation during the day, respectively. The key CAM enzyme PEPC kinase (PPCK; down-regulated during the day) is highlighted in blue font. Species designations are Taust, T. australis - C3; Tspha, T.sphaerocephala – CAM; Tfasc, T. fasciculata – CAM; Tflor, T. floribunda - CAM. Color labels for species are consistent with figures relating to RNA-seq results.

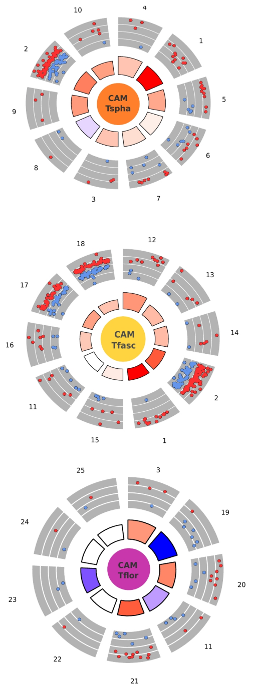


| No. | Species | | | ID | Description |
|---|---|---|---|---|---|
| 1 | * | * | * | GO:0006096 | glycolytic process |
| 2 | * | * | * | GO:0006796 | phosphate–containing compound metabolic ... |
| 3 | * | √ | * | GO:0015977 | carbon fixation |
| 4 | * | √ | √ | GO:1902358 | sulfate transmembrane transport |
| 5 | * | * | * | GO:0015992 | proton transport |
| 6 | * | √ | √ | GO:0006457 | protein folding |
| 7 | * | √ | √ | GO:0006278 | RNA–dependent DNA replication |
| 8 | * | √ | √ | GO:0019430 | removal of superoxide radicals |
| 9 | * | √ | √ | GO:0010019 | chloroplast–nucleus signaling pathway |
| 10 | * | √ | √ | GO:0045489 | pectin biosynthetic process |
| 11 | √ | * | * | GO:0034599 | cellular response to oxidative stress |
| 12 | √ | * | * | GO:0000272 | polysaccharide catabolic process |
| 13 | * | * | * | GO:0006662 | glycerol ether metabolic process |
| 14 | √ | * | * | GO:0000103 | sulfate assimilation |
| 15 | √ | * | * | GO:0009738 | abscisic acid–activated signaling pathwa... |
| 16 | √ | * | * | GO:0045454 | cell redox homeostasis |
| 17 | √ | * | * | GO:0006811 | ion transport |
| 18 | * | * | * | GO:0055085 | transmembrane transport |
| 19 | √ | √ | * | GO:0000280 | nuclear division |
| 20 | √ | √ | * | GO:0032259 | methylation |
| 21 | √ | √ | * | GO:0006820 | anion transport |
| 22 | √ | √ | * | GO:0006383 | transcription from RNA polymerase III pr... |
| 23 | X | √ | * | GO:0050982 | detection of mechanical stimulus |
| 24 | √ | X | * | GO:0006487 | protein N–linked glycosylation |
| 25 | √ | √ | * | GO:0006651 | diacylglycerol biosynthetic process |

NIGHT

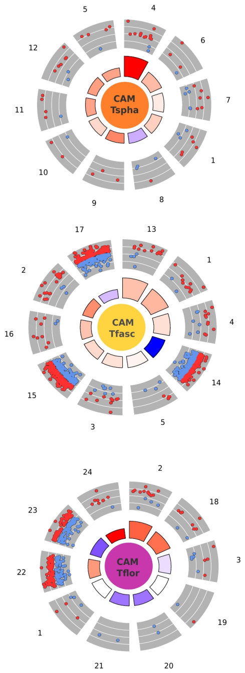


| No. | Species | | | ID | Description |
|---|---|---|---|---|---|
| 1 | * | * | * | GO:0006662 | glycerol ether metabolic process |
| 2 | * | * | * | GO:0006096 | glycolytic process |
| 3 | * | * | * | GO:0034599 | cellular response to oxidative stress |
| 4 | * | * | √ | GO:0006099 | tricarboxylic acid cycle |
| 5 | * | * | √ | GO:0006094 | gluconeogenesis |
| 6 | * | X | √ | GO:0046417 | chorismate metabolic process |
| 7 | * | √ | * | GO:0018105 | peptidyl–serine phosphorylation |
| 8 | * | √ | √ | GO:0070932 | histone H3 deacetylation |
| 9 | * | √ | * | GO:0010019 | chloroplast–nucleus signaling pathway |
| 10 | * | X | √ | GO:0006564 | L–serine biosynthetic process |
| 11 | * | * | X | GO:0009817 | defense response to fungus, incompatible... |
| 12 | * | * | * | GO:0000103 | sulfate assimilation |
| 13 | √ | * | √ | GO:0009735 | response to cytokinin |
| 14 | √ | * | √ | GO:0090304 | nucleic acid metabolic process |
| 15 | √ | * | * | GO:0006796 | phosphate–containing compound metabolic ... |
| 16 | * | * | * | GO:1903959 | regulation of anion transmembrane transp... |
| 17 | * | * | √ | GO:0043412 | macromolecule modification |
| 18 | * | √ | * | GO:0015991 | ATP hydrolysis coupled proton transport |
| 19 | √ | √ | * | GO:1903826 | arginine transmembrane transport |
| 20 | * | * | * | GO:0009870 | defense response signaling pathway, resi... |
| 21 | X | * | * | GO:0050982 | detection of mechanical stimulus |
| 22 | √ | √ | * | GO:0006810 | transport |
| 23 | * | √ | * | GO:0009059 | macromolecule biosynthetic process |
| 24 | √ | √ | * | GO:0009060 | aerobic respiration |

* : significant (FDR 5%)
√ : present in the GO enrichment however not signigificantly (pval > 0.05)
X : not present in the GO enirchment analysis

logFC: downregulated, upregulated
z–score: decreasing, increasing

**SI_Figure_5:** Gene ontology (GO) enrichment results for interspecific CAM-C3 comparisons Results of gene ontology (GO) enrichment analysis for significant genes (FDR<0.05) from transcriptome-wide "interspecific day/night" DE tests. Shown are the 10 most significantly enriched GO terms for each species, excluding GO terms represented by less than two genes. The two subfigures entitled „DAY" and „NIGHT" are composed by the same graphical elements. Subfigure, Right: Rosette plots for genes within each of the enriched GO terms, with red and blue dots indicating up- and down-regulation of single genes, respectively. Wedges in the inner portions of the rosettes designate z-scores based on logFC values for each gene in each group, thus indicating general trends of up- or down- expression in each group (red, increasing: blue, decreasing). Subfigure, Left: Identities and descriptions of the top GO terms, including their occurrence in each species. *: significant enrichment at the 5% level; check-symbol: GO term found in a particular species but no significant enrichment; X: GO term not found in a particular species. Colored cells in the table (right) correspond to those GO terms depicted in the rosette plots (left).

Carbon fixation (ko00710)

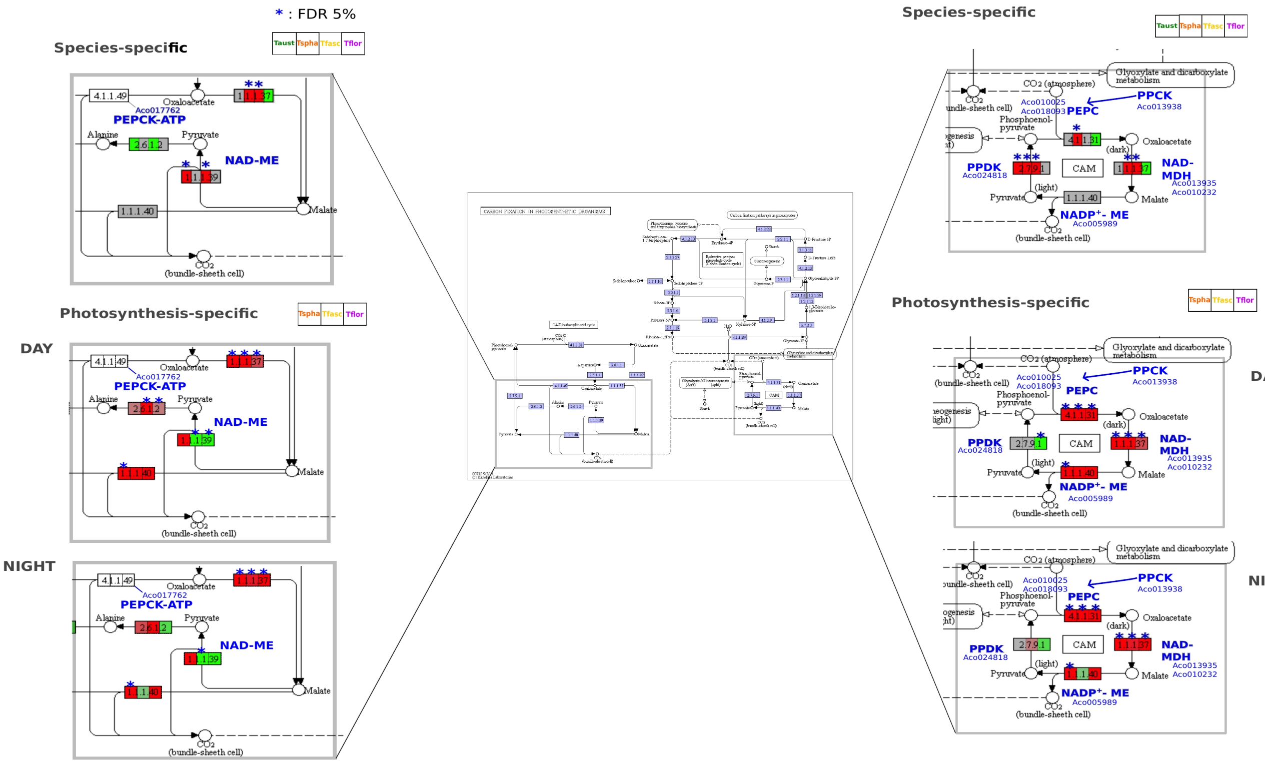


**SI_Figure_6A:** KEGG carbon fixation (ko00710)

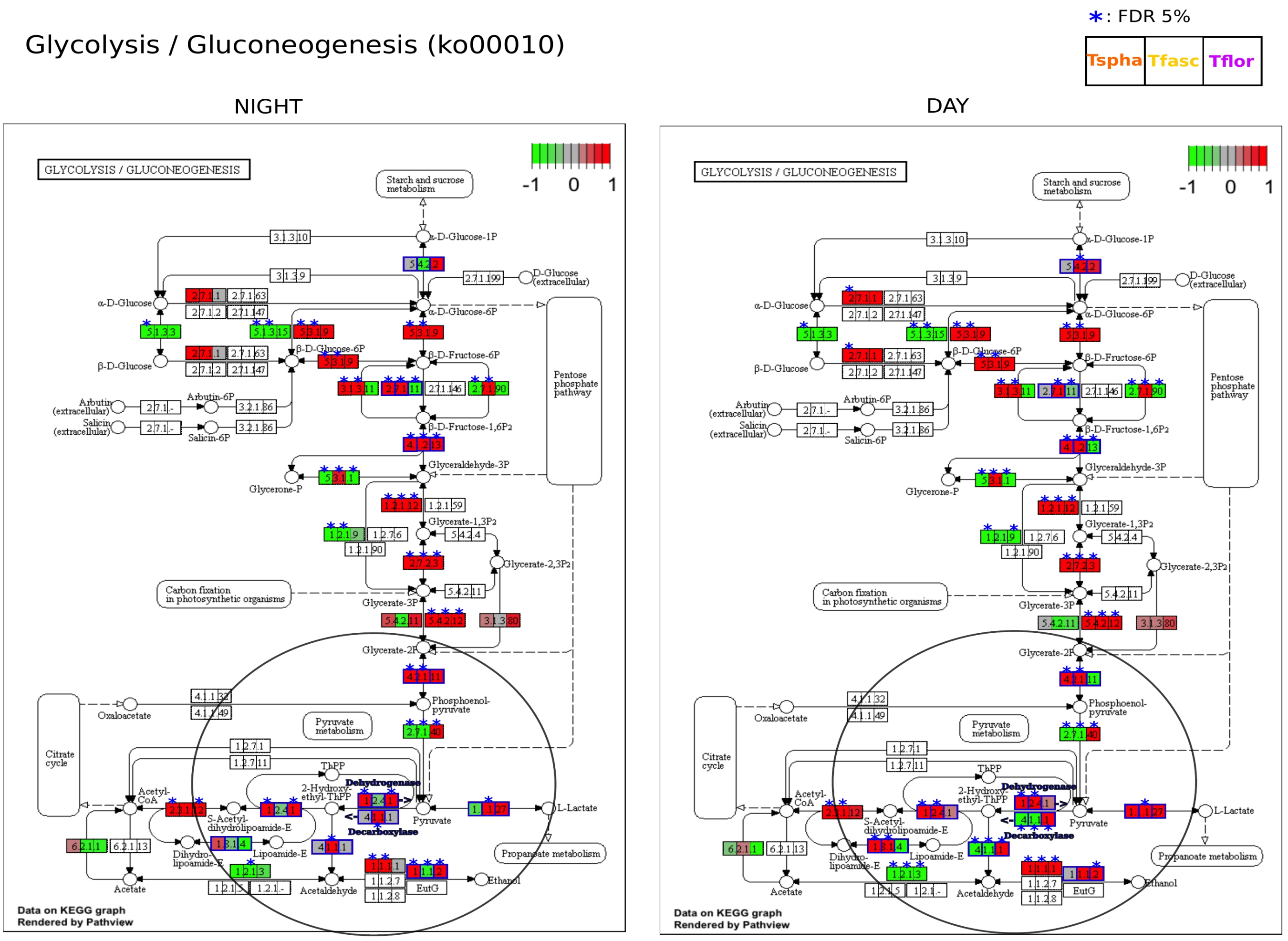


**SI_Figure_6B:** KEGG Starch/sucrose (ko00500)

#### Starch / sucrose (ko00500)

*: FDR 5%

| Tspha | Tfasc | Tflor |
|---|---|---|

#### NIGHT

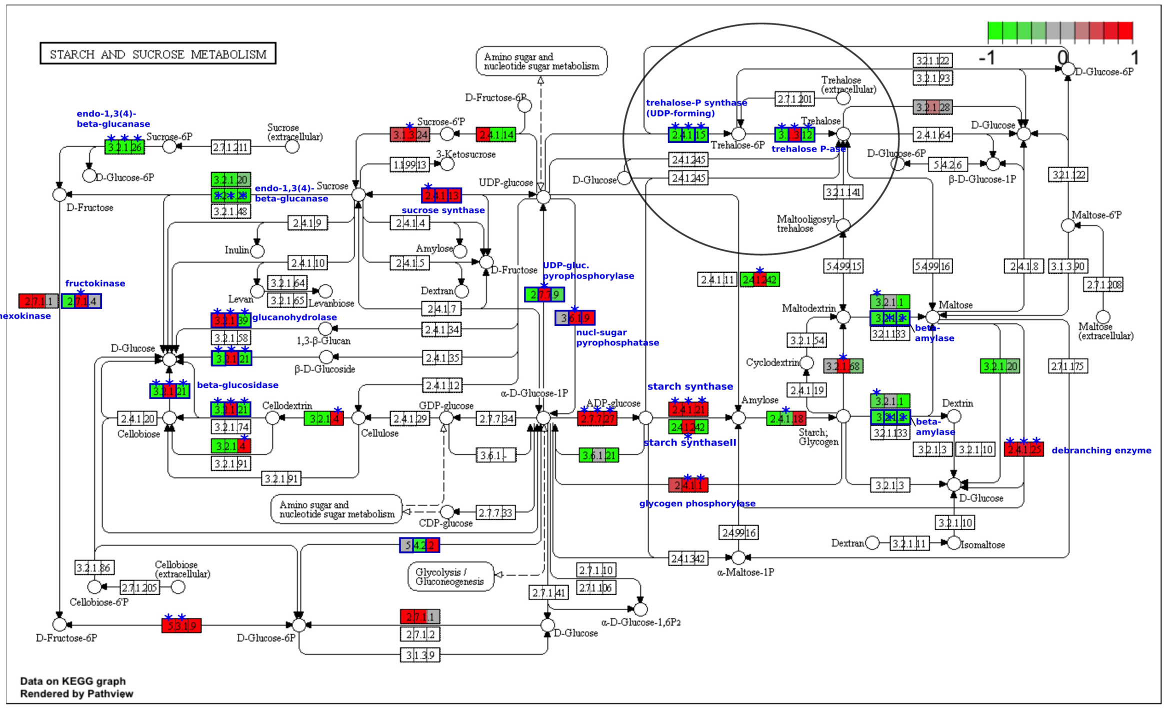


#### DAY

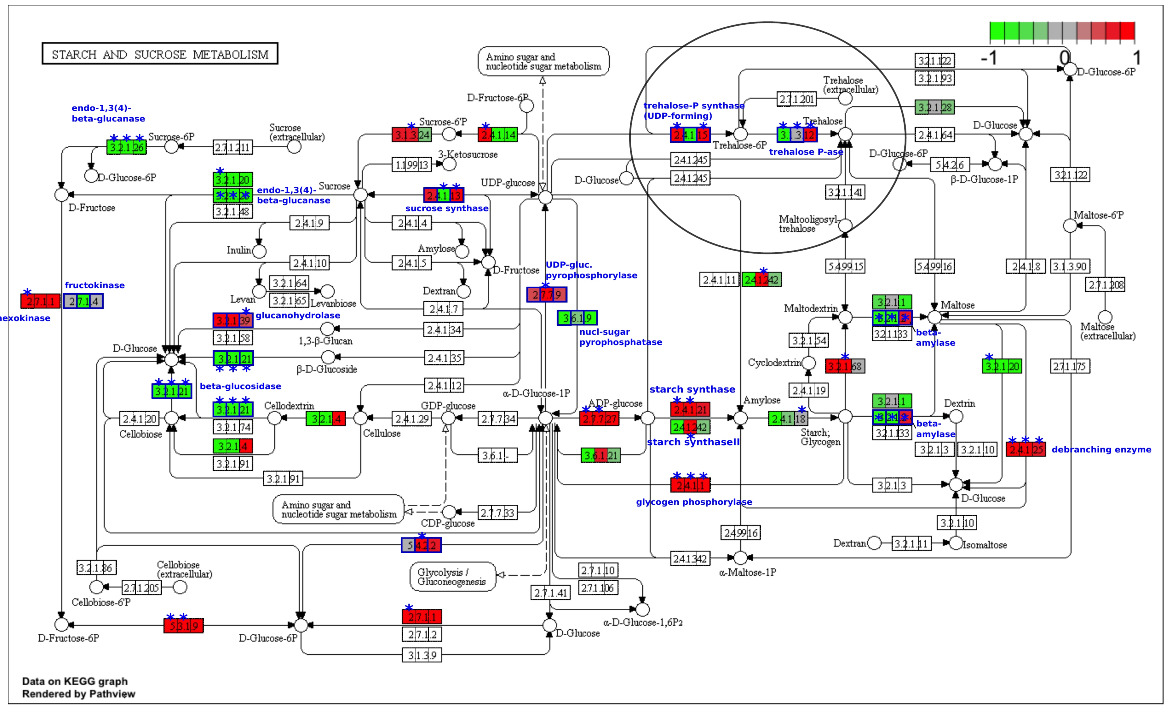


**SI_Figure_6C:** KEGG Glycolysis/gluconeogenesis (ko00010)

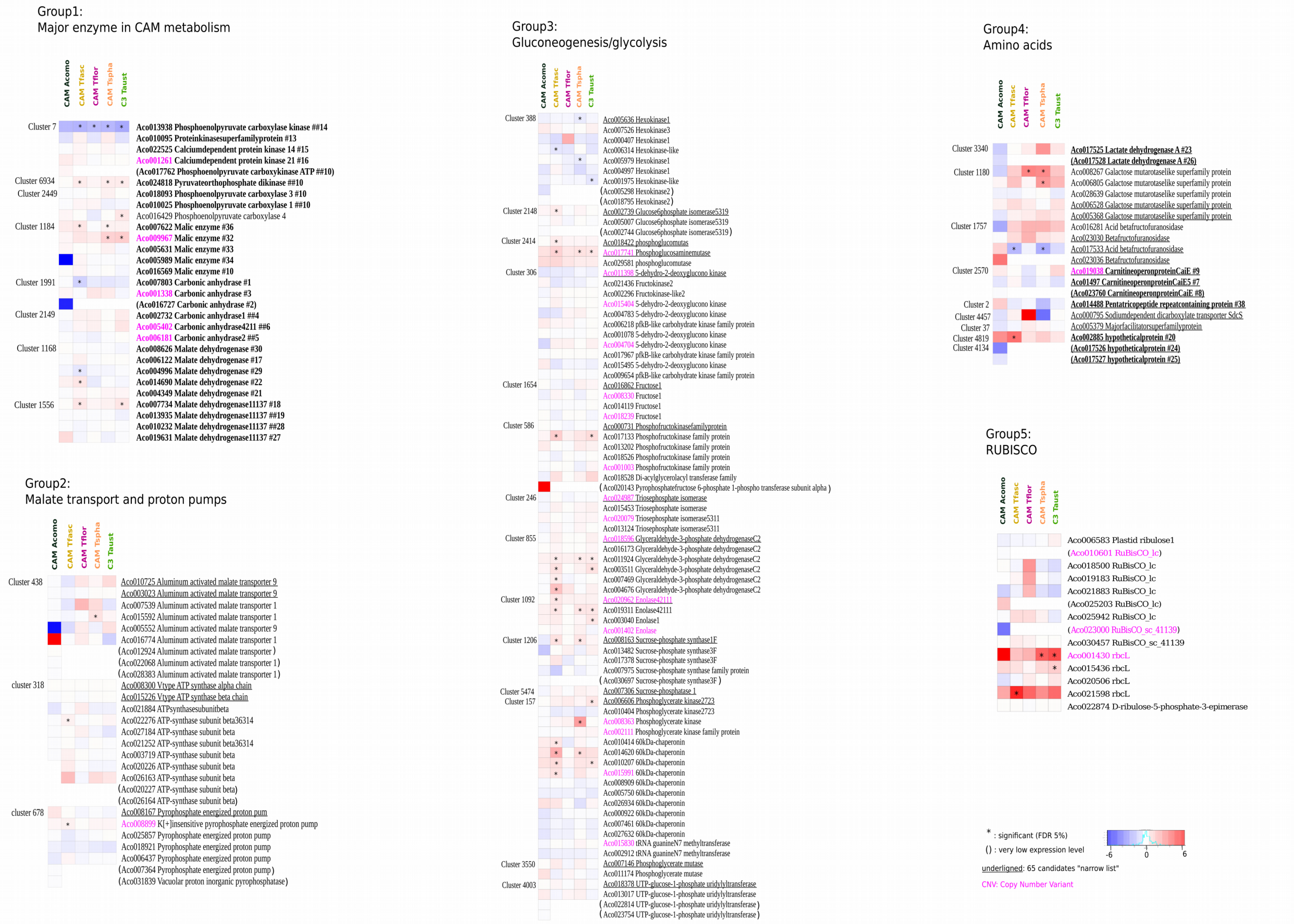

**SI_Figure_7:** Extended target gene list intraspecific test with Pineapple expression data Heatmaps depicting intraspecific day/night DE patterns for homologue clusters of genes with potential involvement in CAM photosynthesis and carbohydrate metabolism based on evidence from pineapple, Ananas comosus (Ming et al. 2015; Wai et al. 2017). Of special relevance to this study are genes involved in gluconeogenesis and glycolysis. The original 65 CAM-related candidate genes used to identify the homologue clusters are underlined.

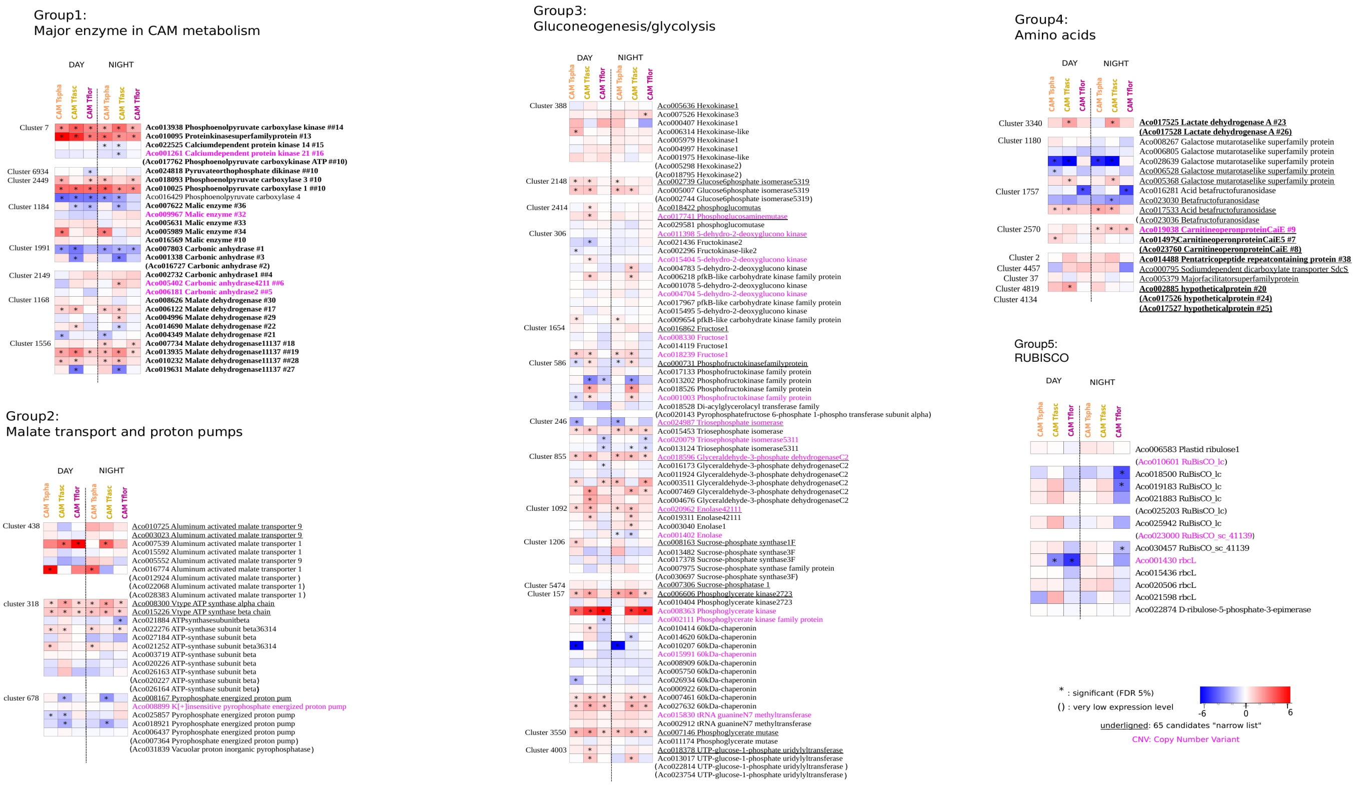


**SI_Figure_8:** Extended target gene list interspecific C3/CAM with Pineapple expression data Heatmaps depicting inter-specific C3/CAM DE patterns for homologue clusters of genes with potential involvement in CAM photosynthesis and carbohydrates metabolism based on evidence from pineapple, *Ananas comosus* (Ming et al. 2015; Wai et al. 2017). Of special relevance are genes involved in gluconeogenesis and glycolysis. The original 65 CAM-related candidate genes used to identify the homologue clusters are underlined.

**SI Tables**

**SI_Table_1:** Data table (DNA-seq and RNA-seq samples table with morphological characters)
**SI_Table_2:** Table of the metabolite analysis (amino acids, carbohydrates and organic acids)
**SI_Table_3:** Positive Selection Gene list
**SI_Table_4: EXCEL** Table of the DE transcripts (233 during the day and 336 during the night) for the interspecific C3/CAM comparisons.
**SI_Table_5: EXCEL** data table (full-genes list with CNV, DE analysis and positive selection test outputs)

### SI Table 1

List of all studied specimens and taxa including essential biological attributes

| Analysis | | | | Samples | Photosynthesis | | Leaves | | | Biological and Ecological characteristics | | | | | | |
|---|---|---|---|---|---|---|---|---|---|---|---|---|---|---|---|---|
| Metabolites | RNA | WGS | WGS mean depth coverage | Species | Photosynthesis | $\Delta^{13}C$ (‰) | Succulence | Shape | Observed_trichomes | Water_tank | Life habit | Bracts | Flower | Pollinators | O: outcrosser S: Self-compatible | Species_geographic range |
| 1 | 0 | 0 | na | Alcantarea regina | C3 | -29.7 | 0 | wide and flat, very fibrous | 0 | 1 | terrestrial | red | yellow | bats | o | Up to 1000m , Brazil |
| 0 | 0 | 1 | 23.8223 | Alcantarea trepida | C3 | -24.6 | 0 | wide and flat | 0 | 1 | saxicolous | brownish /reddish | yellowish/braun | na | o | 300–900 m, Brazil |
| 1 | 0 | 1 | 5.23274 | Barfussia laxissima | C3 | -26.2 | 0 | wide and flat, very fibrous | 0 | 1 | terrestrial | purple | White-pinkish | | o | Bolivia |
| 1 | 0 | 1 | 5.33546 | Lemeltonia dodsonii | C3 | -24.8 | 0 | elongated,thin, delicate | 0 | 0 | epiphytic | green | white | na | o | 1100 m, Ecuador |
| 1 | 0 | 1 | 6.5288 | Pseudalcantarea macropetala | C3 | -31.7 | 0 | wide and flat | 0 | 1 | epiphyte in forest | green | white-yellowish | bats | S/O | Mexico |
| 0 | 0 | 1 | 4.78661 | Racinaea spiculosa | C3 | -25.8 | 0 | wide and flat | 0 | 0 | epiphyte in forest | green | white | na | S/O | Wide south AM distribution |
| 1 | 0 | 0 | na | Racinea sinuosa | C3 | -26.5 | 0 | wide and flat | 1 | 1 | epiphyte | pale yellow | pale yellow | insects | s | na |
| 1 | 0 | 0 | na | T. lajensis | CAM | -20.0 | 0 | wide and flat | 0 | 0 | terrestrial | stems are red and bracts are yellow | pink | na | S/O | na |
| 1 | 0 | 0 | na | T.stricta var albifolia | CAM | -15.4 | 1 | very thick cuticle | 1 | 0 | Epiphytic in dry or wet forest | pink | blue | na | S/O | Brazil |
| 0 | 0 | 1 | 50.4868 | Tillandsia adpressiflora | C3 | -29.3 | 0 | wide and flat | 0 | 0 | Saxicolous or epiphytic | red | dark purple | na | O | 100-625 m, wide South AM distribution |
| 0 | 1 | 1 | 6.29877 | Tillandsia aff. australis | C3 | -27.3 | 0 | wide and flat, very fibreous | 0 | 1 | Saxicolous and epiphytic | red | purple | na | O | 700-3900 m, Bolivia, Argentina |
| 1 | 0 | 0 | na | Tillandsia aff. divaricata | CAM | -15.2 | unclear | wide, flat | 1 | 0 | epiphyte | yellowish/pink | lilac | insect/humming bird | O | na |
| 1 | 1 | 1 | 22.8587 | Tillandsia australis | C3 | -29.0 | 0 | wide and flat, very fibreous | 0 | 1 | Saxicolous and epiphytic | red | purple | na | O | 700-3900 m, Bolivia, Argentina |
| 1 | 1 | 1 | 22.8587 | Tillandsia australis | C3 | -29.0 | 0 | wide and flat, very fibreous | 0 | 1 | Saxicolous and epiphytic | red | purple | na | O | 700-3900 m, Bolivia, Argentina |
| 0 | 1 | 0 | na | Tillandsia australis | C3 | -27.6 | 0 | wide and flat, very fibreous | 0 | 1 | Saxicolous and epiphytic | red | purple | na | O | 700-3900 m, Bolivia, Argentina |
| 1 | 1 | 0 | na | Tillandsia australis | C3 | -27.6 | 0 | wide and flat, very fibreous | 0 | 1 | Saxicolous and epiphytic | red | purple | na | O | 700-3900 m, Bolivia, Argentina |
| 0 | 1 | 0 | na | Tillandsia australis | C3 | -26.1 | 0 | wide and flat, very fibreous | 0 | 1 | Saxicolous and epiphytic | red | purple | na | O | 700-3900 m, Bolivia, Argentina |

**SI Table 1** (continued)

| Analysis | | | | Samples | Photosynthesis | | Leaves | | | Biological and Ecological characteristics | | | | | | |
|---|---|---|---|---|---|---|---|---|---|---|---|---|---|---|---|---|
| Metabolites | RNA | WGS | WGS mean depth coverage | Species | Photosynthesis | $\Delta^{13}C$ (‰) | Succulence | Shape | Observed_trichomes | Water_tank | Life habit | Bracts | Flower | Pollinators | O: outcrosser S: Self-compatible | Species_geographic range |
| 1 | 0 | 0 | na | Tillandsia capillaris | CAM | -15.0 | 1 | curly, very tinny | 1 | 0 | epiphyte | yellow | green | na | S/O | na |
| 1 | 0 | 0 | na | Tillandsia complanata | C3 | -30.9 | 0 | wide and flat | 1 | 1 | epiphyte in forest | red | pink | hummingbird | O | 750-3600 m broad South AM distribution |
| 0 | 0 | 1 | 7.09465 | Tillandsia curvispica | CAM | -22.9 | 1 | very thick cuticl | 1 | 0 | saxicolous | green | white | na | O | 1200m, Peru |
| 1 | 0 | 1 | 6.07676 | Tillandsia demissa | CAM | -22.0 | 0 | wide and flat, not fibrous | 0 | 1 | Saxicolous and epiphytic | yellowish/pink | lilac/pinkish | na | O | Ecuador, Loja |
| 1 | 0 | 1 | 6.03045 | Tillandsia disticha | CAM | -17.6 | 1 | elongated,thin | 1 | 0 | epiphyte | green | yellow | na | O | Colombia, Ecuador, Peru |
| 0 | 0 | 1 | 5.71611 | Tillandsia divaricata | CAM | -15.8 | 1 | thick cuticle | 1 | 0 | epiphyte | red | pink, red | na | O | Ecuador |
| 0 | 1 | 1 | 30.0668 | Tillandsia fasciculata | CAM | -15.8 | 1 | very thick cuticle | 1 | 0 | Epiphytic in woods | Yellow and red | purple | na | s | 0 to 1350m, Central and South AM, West Indies |
| 0 | 1 | 0 | na | Tillandsia fasciculata | CAM | -18.3 | 1 | very thick cuticle | 1 | 0 | Epiphytic in woods | Yellow and red | purple | na | s | 0 to 1350m, Central and South AM, West Indies |
| 1 | 1 | 0 | na | Tillandsia fasciculata | CAM | -18.3 | 1 | very thick cuticle | 1 | 0 | Epiphytic in woods | Yellow and red | purple | na | s | 0 to 1350m, Central and South AM, West Indies |
| 0 | 1 | 0 | na | Tillandsia fasciculata | CAM | -16.1 | 1 | very thick cuticle | 1 | 0 | Epiphytic in woods | Yellow and red | purple | na | S/O | 0 to 1350m, Central and South AM, West Indies |
| 1 | 1 | 0 | na | Tillandsia fasciculata | CAM | -16.1 | 1 | very thick cuticle | 1 | 0 | Epiphytic in woods | Yellow and red | purple | na | s | 0 to 1350m, Central and South AM, West Indies |
| 0 | 1 | 0 | na | Tillandsia fasciculata | CAM | -14.5 | 1 | very thick cuticle | 1 | 0 | Epiphytic in woods | Yellow and red | purple | na | s | 0 to 1350m, Central and South AM, West Indies |
| 1 | 1 | 0 | na | Tillandsia fasciculata | CAM | -14.5 | 1 | very thick cuticle | 1 | 0 | Epiphytic in woods | Yellow and red | purple | na | s | 0 to 1350m, Central and South AM, West Indies |
| 0 | 1 | 0 | na | Tillandsia floribunda | CAM | -18.6 | 1 | elongated,thin | 1 | 0 | Saxicolous and epiphytic | red | purple | na | O | 900-2500 m, Ecuador – Peru |
| 1 | 1 | 1 | 26.4765 | Tillandsia floribunda | CAM | -18.6 | 1 | elongated,thin | 1 | 0 | Saxicolous and epiphytic | red | purple | na | O | 900-2500 m, Ecuador – Peru |
| 0 | 1 | 0 | na | Tillandsia floribunda | CAM | -20.3 | 1 | elongated,thin | 1 | 0 | Saxicolous and epiphytic | red | purple | na | O | 900-2500 m, Ecuador – Peru |
| 1 | 1 | 0 | na | Tillandsia floribunda | CAM | -20.3 | 1 | elongated,thin | 1 | 0 | Saxicolous and epiphytic | red | purple | na | O | 900-2500 m, Ecuador – Peru |

**SI Table 1** (continued)

| Analysis | | | | Samples | Photosynthesis | | Leaves | | | Biological and Ecological characteristics | | | | | | |
|---|---|---|---|---|---|---|---|---|---|---|---|---|---|---|---|---|
| Metabolites | RNA | WGS | WGS mean depth coverage | Species | Photosynthesis | $\Delta^{13}C$ (‰) | Succulence | Shape | Observed_trichomes | Water_tank | Life habit | Bracts | Flower | Pollinators | O: outcrosser S: Self-compatible | Species_geographic range |
| 1 | 1 | 0 | na | Tillandsia floribunda | CAM | -18.9 | 1 | elongated,thin | 1 | 0 | Saxicolous and epiphytic | red | purple | na | O | 900-2500 m, Ecuador – Peru |
| 0 | 0 | 1 | na | Tillandsia gardneri | CAM | -19.36 | 1 | very thick cuticle | 1 | 0 | Epiphytic and saxicolous | pink | pink/red | na | S/O | Wide south AM distribution |
| 0 | 0 | 1 | 23.1307 | Tillandsia hitchcockiana | CAM | -17.6 | unclear | very thick cuticle | 1 | 0 | epiphyte | red | pinkish | na | O | 1200--1900 m, Ecuador, Peru |
| 1 | 0 | 0 | na | Tillandsia hitchcockiana | CAM | -17.32 | 1 | very thick cuticle | 1 | 0 | epiphyte | red | pinkish | na | O | 1200--1900 m, Ecuador, Peru |
| 1 | 0 | 0 | na | Tillandsia juncea | CAM | -15.35 | 0 | elongated,thin | 1 | 0 | Terrestrial to epiphyte | red | purple | na | S/O | 5-2416 m, Wide south AM distribution |
| 0 | 0 | 1 | 20.6196 | Tillandsia juncea | CAM | -21.6 | 0 | elongated,thin | 1 | 0 | Terrestrial to epiphyte | red | purple | na | S/O | 5-2416 m, Wide south AM distribution |
| 1 | 0 | 0 | na | Tillandsia latifolia | CAM | -18.1 | 1 | flat and curly | 1 (a lot) | 0 | epiphyte | red | pink | na | S/O | na |
| 0 | 0 | 1 | 21.0746 | Tillandsia latifolia latifolia | CAM | -18.5 | 1 | thick cuticle | 1 | 0 | saxicolous | red | pink | na | O | 0 to 2900 m, Peru |
| 1 | 0 | 1 | 13.2354 | Tillandsia leiboldiana | C3 | -31.3 | 0 | wide and flat | 0 | 1 | epiphyte in forest | red | purple | na | O | 25-2000 m, Central AM |
| 1 | 0 | 1 | 29.6024 | Tillandsia propagulifera | CAM | -20.4 | 1 | thin | 1 | 0 | epiphytic | green | deep purple to black | small moths | O | 450 m, Peru |
| 1 | 0 | 1 | 6.13085 | Tillandsia rauhii | CAM | -20.7 | 0 | Flat, very fibrous | 1 | 1 | saxicolous | green | deep purple to black | na | O | 700 m, Central AM and Peru |
| 0 | 0 | 1 | 30.4144 | Tillandsia somnians | C3 | -26.9 | 0 | wide and flat, not fibrous | 0 | 1 | saxicolous | red | purple | na | O | 600m, Ecuador - Peru |
| 1 | 0 | 0 | na | Tillandsia somnians | C3 | -26.9 | 0 | wide and flat, not fibrous | 0 | 1 | saxicolous | red | purple | na | O | 600m, Ecuador - Peru |
| 1 | 1 | 1 | 26.227 | Tillandsia sphaerocephala | CAM | -15.7 | unclear | wide and flat | 1 | 0 | Saxicolous and epiphytic | red | purple | na | O | Peru-Bolivia-Argentina |
| 0 | 1 | 0 | na | Tillandsia sphaerocephala | CAM | -16.2 | unclear | wide and flat | 1 | 0 | Saxicolous and epiphytic | red | purple | na | O | Peru-Bolivia-Argentina |
| 1 | 1 | 0 | na | Tillandsia sphaerocephala | CAM | -16.2 | unclear | wide and flat | 1 | 0 | Saxicolous and epiphytic | red | purple | na | O | Peru-Bolivia-Argentina |
| 0 | 1 | 0 | na | Tillandsia sphaerocephala | CAM | -15.7 | unclear | wide and flat | 1 | 0 | Saxicolous and epiphytic | red | purple | na | O | Peru-Bolivia-Argentina |
| 1 | 1 | 0 | na | Tillandsia sphaerocephala | CAM | -15.7 | unclear | wide and flat | 1 | 0 | Saxicolous and epiphytic | red | purple | na | O | Peru-Bolivia-Argentina |
| 0 | 1 | 0 | na | Tillandsia sphaerocephala | CAM | -15.7 | unclear | wide and flat | 1 | 0 | Saxicolous and epiphytic | red | purple | na | O | Peru-Bolivia-Argentina |
| 0 | 0 | 1 | 25.9628 | Tillandsia stenoura | C3 | -25.7 | 0 | wide and flat | 0 | 1 | terrestrial | red | purple | na | O | na |
| 0 | 0 | 1 | na | Tillandsia stricta | CAM | -16.5 | 1 | very thick cuticle | 1 | 0 | Epiphytic | pink | blue | na | O | Wide south AM distribution |

**SI Table 1** (continued)

| Analysis | | | | Samples | Photosynthesis | | Leaves | | | Biological and Ecological characteristics | | | | | | |
|---|---|---|---|---|---|---|---|---|---|---|---|---|---|---|---|---|
| Metabolites | RNA | WGS | WGS mean depth coverage | Species | Photosynthesis | $\Delta^{13}C$ (‰) | Succulence | Shape | Observed_trichomes | Water_tank | Life habit | Bracts | Flower | Pollinators | O: outcrosser S: Self-compatible | Species_geographic range |
| 0 | 0 | 1 | 23.8212 | Tillandsia trauneri | C3 | -25.7 | 0 | wide and flat, very fibrous | 0 | 1 | Saxicolous | red | dark purple, almost black | na | O | 1100m, Mexico |
| 0 | 0 | 1 | 14.7596 | Vriesea itatiaiae | C3 | -25.56 | 0 | wide and flat | 0 | 1 | epiphyte and saxicolous | yellow | red | bat | O | 2000-2300 m, Brazil |
| 1 | 0 | 0 | na | Vriesea longicaulis | C3 | -26.1 | 0 | wide and flat | 0 | 1 | terrestrial, epiphyte in forest | green | brownish | bats | O | 750- 1900 m, Brazil |
| 1 | 0 | 1 | 4.93188 | Wallisia lindeniana | C3 | -26.9 | 0 | wide and flat | 0 | 0 | epiphyte | green | from white to blue/purple | na | O | na |
| 0 | 0 | 1 | 26.3262 | na | na | na | na | na | na | na | na | na | na | na | na | na |

**SI_Table_2:** Abundance data for compounds identified by targeted metabolite analysis (1/6)

| species | sampling_time | isotope | [illegible] | Citric_acid | Malic_acid | Fructose_methoxyamine | Fructose_methoxyamine_BP | Glucose_MP | Glucose_methoxyamine_BP | Sucrose | myo_Inositol | Alanine | Asparagine | Aspartic_acid | Fumaric_acid | Galactinol | Galactose_methoxyamine | Galactose_methoxyamine_BP | Glutamic_acid | GlutamineDL | Glutaric_acid_2oxo_BP |
|---|---|---|---|---|---|---|---|---|---|---|---|---|---|---|---|---|---|---|---|---|---|
| *Alcantarea_regina** | day | -29.7 | C3 | 3579.69 | 11590.3 | 232465.1691 | 147867 | 98824.6 | 11176.3 | 268325 | 88166.1 | 118294 | 1226183 | 331697 | 3516.14 | 15778.3 | 17681.8 | 243821 | 242448 | 2131465 | 10029.1 |
| *Alcantarea_regina** | night | -29.7 | C3 | 8413.73 | 26669 | 396149.6437 | 265354 | 229033 | 32637.5 | 320714 | 261531 | 0 | 245325 | 65336.5 | 40015.2 | 18660.8 | 87234.6 | 50893.6 | 115682 | 463817 | 98970.8 |
| *Barfussia_laxissima* | night | -26.2 | C3 | 6967.67 | 825.036 | 33975.39192 | 22774.7 | 3483.8 | 386.366 | 106260 | 1540.64 | 71300 | 29496.6 | 37172.3 | 0 | 55221.9 | 136.221 | 13025.2 | 57977.3 | 29380.8 | 3415.72 |
| *Barfussia_laxissima* | day | -26.2 | C3 | 4767.06 | 1375.41 | 155987.7454 | 109737 | 28482.8 | 3092.84 | 219130 | 2372.57 | 127509 | 7676.21 | 35990.6 | 0 | 28440.7 | 12308.3 | 34434.8 | 29822.5 | 24853.7 | 4383.74 |
| *Tillandsia_australis* | night | -29.0 | C3 | 16285.7 | 22222.1 | 67933.12195 | 41015.6 | 332773 | 69296 | 275010 | 65799.8 | 147933 | 2503503 | 785564 | 900.805 | 4909.39 | 87599.9 | 64954.1 | 128487 | 502877 | 37059.5 |
| *Tillandsia_australis* | day | -29.0 | C3 | 12337 | 45428.7 | 326512.4163 | 196790 | 198249 | 23829.3 | 130831 | 28662.1 | 49866.5 | 154113 | 43859 | 0 | 2169.04 | 31714.4 | 22046.1 | 19304.1 | 9122.73 | 7618.04 |
| *Tillandsia_leiboldiana* | night | -31.3 | C3 | 56452.8 | 1568.42 | 701774.6891 | 495472 | 359989 | 78465.8 | 396287 | 51919.9 | 18175.5 | 8006.14 | 11044.3 | 2112.18 | 16091.1 | 91222.4 | 103676 | 15526.4 | 3444.74 | 3222.69 |
| *Tillandsia_leiboldiana* | day | -31.3 | C3 | 20604.7 | 5888.64 | 710944.4976 | 529729 | 311476 | 62454.8 | 472195 | 82947.5 | 55198.5 | 1082.18 | 30551.3 | 438.708 | 38510.1 | 81959.9 | 48318.7 | 54247.9 | 3132.25 | 9322.87 |
| *Tillandsia_propagulifera* | night | -20.4 | CAM | 55003.6 | 134486 | 4316.828645 | 2544.88 | 12650.5 | 1376.65 | 230410 | 81528.9 | 29025 | 1094524 | 189998 | 25970.4 | 2195.55 | 769.156 | 46905.5 | 45559.5 | 29601 | 18435.5 |
| *Tillandsia_propagulifera* | day | -20.4 | CAM | 78657.8 | 163979 | 6449.868074 | 3751.87 | 26077.8 | 2854.01 | 180135 | 115705 | 25195.2 | 1200957 | 224245 | 288589 | 4878.13 | 113.596 | 4033.35 | 70696.7 | 65227.1 | 41121.3 |
| *Tillandsia_floribunda* | night | -18.6 | CAM | 241355 | 79520.8 | 54265.22459 | 34154.8 | 43658 | 5073.88 | 211129 | 33986.2 | 267903 | 1921024 | 392970 | 9814 | 19172 | 7256.24 | 0 | 66540.1 | 223276 | 16934.7 |
| *Tillandsia_floribunda* | day | -18.6 | CAM | 216518 | 75194.3 | 61624.86413 | 38391.8 | 75342.2 | 9126.9 | 227369 | 33070.3 | 218316 | 2446024 | 344985 | 9846.39 | 22539.6 | 6395.24 | 31198.6 | 95012 | 353192 | 18908.5 |
| *Tillandsia_latifolia_ssp_* | night | -18.1 | CAM | 139083 | 68411.2 | 71666.3 | 45106.2 | 6060.72 | 528.082 | 114985 | 367466 | 53678.2 | 150056 | 72590 | 3605.86 | 15889.2 | 2095.7 | 18496.6 | 24422.9 | 70.9591 | 1482.92 |
| *Tillandsia_latifolia_ssp_* | day | -18.1 | CAM | 45142.6 | 10957.7 | 1071684.5 | 837742 | 373925 | 98379.4 | 273024 | 383799 | 18908.7 | 4512.4 | 14840.9 | 292.398 | 19036.1 | 38.952 | 94438.3 | 13474.1 | 1619.21 | 9804.7 |
| *Vriesea_hitchcockiana* | night | -17.32 | CAM | 51636.4 | 151419 | 485943.9803 | 323370 | 75260.6 | 7665.45 | 93876.5 | 74266.6 | 87255.9 | 393415 | 271639 | 15822.7 | 6020.98 | 4726.31 | 26561.9 | 41460.5 | 231699 | 13309.2 |
| *Vriesea_hitchcockiana* | day | -17.32 | CAM | 27648.1 | 140420 | 843775.6675 | 579679 | 150704 | 16817.5 | 110547 | 63889.6 | 44473.7 | 152878 | 76529.6 | 2317.63 | 5398.77 | 3527.51 | 21397 | 24343.5 | 29838.3 | 9696.9 |
| *Tillandsia_sphaerocepha* | night | -15.7 | CAM | 44877.3 | 40909.3 | 210268.1343 | 148540 | 39280.1 | 4203.06 | 66242.6 | 106031 | 43453.6 | 3501359 | 534986 | 2072.84 | 26410.9 | 10784.3 | 82391.2 | 50767.2 | 70413.3 | 6905.12 |
| *Tillandsia_sphaerocepha* | day | -15.7 | CAM | 94770.3 | 123980 | 263305.75 | 158974 | 38310.4 | 3757.94 | 109041 | 295168 | 27017 | 315595 | 56531.3 | 3598.25 | 1536.25 | 7045.25 | 0 | 35721.1 | 5499.14 | 9662.67 |
| *Tillandsia_somnians* | night | -26.9 | C3 | 17586 | 9720.79 | 490483.7799 | 313865 | 77409.3 | 8094.16 | 73389.7 | 106807 | 789059 | 63558.9 | 200289 | 35.9282 | 1544.28 | 11376 | 6447.08 | 34182.5 | 667622 | 3960.67 |
| *Tillandsia_somnians* | day | -26.9 | C3 | 22909.1 | 1421.59 | 853008.8806 | 573577 | 334710 | 67739.9 | 249032 | 345374 | 37987.3 | 12864.4 | 19423.6 | 522.711 | 2061.24 | 31763.2 | 27983.7 | 13397.5 | 12804.2 | 9168.78 |
| *Tillandsia_complanata* | night | -30.9 | C3 | 12097 | 0 | 1934.156977 | 1448.81 | 323.837 | 0 | 224486 | 30355.1 | 101197 | 1959322 | 186616 | 0 | 19966.9 | 12.3826 | 1730 | 92029.8 | 57453.7 | 0 |

**SI_Table_2:** Abundance data for compounds identified by targeted metabolite analysis (2/6)

| species | sampling_time | isotope | [illegible] | Citric_acid | Malic_acid | ructose_methoxyamine | ructose_methoxyamine_B | Glucose_MP | lucose_methoxyamine_BI | Sucrose | myo_Inositol | Alanine | Asparagine | Aspartic_acid | Fumaric_acid | Galactinol | ialactose_methoxyamine | ialactose_methoxyamine_ | Glutamic_acid | GlutamineDL | Glutaric_acid_2oxo_BP |
|---|---|---|---|---|---|---|---|---|---|---|---|---|---|---|---|---|---|---|---|---|---|
| *Tillandsia_complanata* | day | -30.9 | C3 | 55309.9 | 4220.5 | 4182.693267 | 2623.04 | 638.978 | 124.017 | 274188 | 41577.3 | 68573 | 2084878 | 161962 | 3.51845 | 43942.7 | 34.0923 | 558.953 | 130299 | 143427 | 849.601 |
| *Lemeltonia_dodsonii* | night | -24.8 | C3 | 252202 | 57726.7 | 14354 | 8816.68 | 135386 | 17969.1 | 249812 | 12138.2 | 123037 | 588691 | 232611 | 5779.77 | 14346.1 | 141.757 | 62082.5 | 71091.2 | 200160 | 3964.35 |
| *Lemeltonia_dodsonii* | day | -24.8 | C3 | 248406 | 17072.8 | 14178.29949 | 8459.06 | 8669.29 | 952.868 | 383714 | 1666.83 | 252288 | 102692 | 779369 | 5538.4 | 15099.2 | 14.2487 | 485.533 | 220281 | 153451 | 3290.69 |
| *Tillandsia_striaca* | night | -15.4 | CAM | 86161.5 | 132859 | 281146.6348 | 196593 | 46089.8 | 5008.93 | 60723.9 | 4560.05 | 27644.5 | 595065 | 149073 | 15374.6 | 3358.54 | 11505.2 | 74758.7 | 52763.8 | 9681.93 | 10781.1 |
| *Tillandsia_striaca* | day | -15.4 | CAM | 166288 | 219644 | 169632.7475 | 114269 | 15848.4 | 1505.07 | 91786.3 | 3736.41 | 19713.8 | 408868 | 94723.6 | 34423.6 | 2684.31 | 3339.26 | 20704.8 | 71229.8 | 1207.52 | 9944.78 |
| *Tillandsia_arcuans(=syn.* | night | -25.7 | C3 | 76414.2 | 23275.9 | 267374.7502 | 171102 | 298186 | 155685 | 264989 | 185685 | 26230.3 | 0 | 32342.8 | 904.57 | 11433.7 | 39618.1 | 56484.3 | 14241.1 | 163.419 | 2513.21 |
| *Tillandsia_arcuans(=syn.* | day | -25.7 | C3 | 94949.5 | 17801 | 217893.1868 | 137415 | 514182 | 198347 | 221603 | 223709 | 21878.6 | 0 | 14573.7 | 1208.08 | 16508.3 | 36545.4 | 32484.9 | 28745.5 | 52.8269 | 3389.26 |
| *Tillandsia_australis°* | night | -26.1 | C3 | 21325.4 | 15380.4 | 23847.06494 | 14217 | 160197 | 22895.1 | 132698 | 191462 | 953639 | 372678 | 668773 | 4711.22 | 5870.7 | 19678.3 | 0 | 15782.6 | 71721.7 | 42557.1 |
| *Tillandsia_australis* | night | -27.6 | C3 | 34145.4 | 32429.8 | 62782.42347 | 37402.7 | 188166 | 24609.3 | 189263 | 81943.9 | 1526667 | 527122 | 690168 | 1688.93 | 11289.4 | 22845.5 | 49352 | 86077.3 | 849371 | 1102.7 |
| *Tillandsia_australis°* | day | -26.1 | C3 | 20154.1 | 19239.2 | 118035.8612 | 73612.4 | 73803.8 | 8875.94 | 98280.2 | 227488 | 2200160 | 175102 | 422723 | 0 | 4583.24 | 7496.58 | 23787.7 | 0 | 30566.7 | 14840.5 |
| *Tillandsia_australis* | day | -27.6 | C3 | 22063.1 | 49074.2 | 45319.65854 | 25971.5 | 352602 | 81858.4 | 264403 | 118391 | 2964589 | 1459635 | 729349 | 595.293 | 17364.7 | 46330.6 | 58776.1 | 81821.8 | 2022642 | 953.22 |
| *Tillandsia_capillaris* | night | -15.0 | CAM | 55261.1 | 89420.7 | 59826.45669 | 37155.1 | 53373.5 | 6107.14 | 18248.3 | 2667.95 | 16434.7 | 1305087 | 142231 | 8471.57 | 0 | 4398.4 | 52411.2 | 39505.7 | 14825.9 | 9222.89 |
| *Tillandsia_capillaris* | day | -15.0 | CAM | 52929.2 | 64671.8 | 37513.02872 | 23342.5 | 42343.9 | 4742.3 | 54889.3 | 2150.57 | 24850.4 | 136955 | 18615.1 | 13397.4 | 0 | 2362.11 | 28136.8 | 27290.9 | 14453.4 | 5974.15 |
| *Tillandsia_fasciculata°* | night | -16.1 | CAM | 45683.6 | 152744 | 1081976.901 | 738638 | 287341 | 47144.8 | 328447 | 396263 | 153798 | 1412450 | 382253 | 55679.2 | 14482.9 | 18964.9 | 123746 | 99751.4 | 256776 | 30874.2 |
| *Tillandsia_fasciculata* | night | -18.3 | CAM | 234249 | 315075 | 34908.11765 | 21231.6 | 28005.8 | 3048.85 | 246287 | 11454.4 | 28378.1 | 1169334 | 166258 | 461104 | 2384.4 | 6532 | 3218.49 | 55113.8 | 6543.18 | 12498.3 |
| *Tillandsia_fasciculata* | night | -14.5 | CAM | 49258.9 | 113308 | 1160.083565 | 713.872 | 78.766 | 0 | 68095.2 | 24730.9 | 28141 | 128996 | 62412.7 | 98993.9 | 6610.39 | 0 | 54.0056 | 38420.2 | 0 | 8261.06 |
| *Tillandsia_fasciculata°* | day | -16.1 | CAM | 87875.8 | 184476 | 326392.3077 | 202385 | 39787.9 | 4038.67 | 240587 | 39484.8 | 79257.1 | 1218675 | 48292.5 | 12480.5 | 8666.07 | 2789.81 | 14044.1 | 49826.4 | 12140.3 | 10404.7 |
| *Tillandsia_fasciculata* | day | -18.3 | CAM | 745654 | 93816.7 | 560696.615 | 342562 | 335336 | 61978.6 | 113947 | 58916.3 | 120281 | 5198285 | 212545 | 122723 | 19707.9 | 27559.7 | 96319.5 | 291678 | 31150.1 | 1258.97 |
| *Tillandsia_fasciculata* | day | -14.5 | CAM | 114511 | 156709 | 166043.4184 | 100835 | 80637.9 | 9021.45 | 293391 | 43209.4 | 59060 | 1976798 | 121324 | 58425.2 | 14088 | 9298.98 | 0 | 72347.3 | 14977 | 13241.6 |
| *Tillandsia_floribunda* | night | -20.3 | CAM | 205730 | 63954 | 19715.6338 | 11739 | 5441.72 | 611.268 | 171670 | 20532.7 | 114544 | 3558901 | 77962.1 | 8741.58 | 4713.27 | 881.521 | 0 | 80041.3 | 197234 | 15261 |
| *Tillandsia_floribunda* | night | -18.9 | CAM | 169003 | 69754.6 | 363101.8293 | 246617 | 89736.9 | 10230.6 | 280403 | 66151 | 310102 | 695726 | 742844 | 3619.15 | 13054.1 | 10032.9 | 52940.9 | 76997 | 390085 | 13108 |
| *Tillandsia_floribunda* | day | -20.3 | CAM | 330291 | 48491 | 26462.5 | 16189.3 | 7897.42 | 910.526 | 121423 | 17720 | 41641.4 | 3322571 | 643502 | 16288.7 | 5793.11 | 2764.63 | 0 | 104670 | 100319 | 10278 |
| *Tillandsia_floribunda* | day | -18.9 | CAM | 169773 | 32681.1 | 588408.8283 | 418006 | 230173 | 29169.9 | 326788 | 91012.8 | 266412 | 669779 | 459338 | 4892.4 | 30444.9 | 22382.6 | 51081.7 | 117823 | 432843 | 579.537 |
| *Tillandsia_sphaerocepha* | night | -16.2 | CAM | 183530 | 82510.8 | 561079.5276 | 371777 | 117577 | 12982.9 | 104528 | 74105 | 38459.8 | 1969020 | 181308 | 4129.87 | 662.835 | 6745.8 | 35846 | 39155.9 | 17123.2 | 7034.59 |
| *Tillandsia_sphaerocepha* | night | -15.7 | CAM | 156274 | 92156.7 | 437431.3178 | 279623 | 86107.8 | 8765.81 | 234357 | 202363 | 58617 | 243738 | 308766 | 230835 | 70.2481 | 9499.38 | 66944.1 | 40526.8 | 37475.7 | 31059.4 |

**SI_Table_2:** Abundance data for compounds identified by targeted metabolite analysis (3/6)

| species | sampling_time | isotope | [illegible] | Citric_acid | Malic_acid | Fructose_methoxyamine | Fructose_methoxyamine_BP | Glucose_MP | Glucose_methoxyamine_BP | Sucrose | myo_Inositol | Alanine | Asparagine | Aspartic_acid | Fumaric_acid | Galactinol | Galactose_methoxyamine | Galactose_methoxyamine_BP | Glutamic_acid | GlutamineDL | Glutaric_acid_2oxo_BP |
|---|---|---|---|---|---|---|---|---|---|---|---|---|---|---|---|---|---|---|---|---|---|
| *Tillandsia_sphaerocepha* | day | -16.2 | CAM | 172105 | 96318.4 | 136544.9448 | 86095.5 | 34832.9 | 3629.61 | 155874 | 143343 | 23387.9 | 65500.2 | 86007.8 | 14324.6 | 8875.55 | 6511.16 | 14884 | 54641.9 | 6395.19 | 14055.8 |
| *Tillandsia_sphaerocepha* | day | -15.7 | CAM | 92095.6 | 9637.33 | 268633.4232 | 159741 | 61975.5 | 6617.71 | 215090 | 268425 | 20757.9 | 42.7951 | 11790.9 | 925.121 | 4039.3 | 6022.88 | 11085.4 | 15348.2 | 371.159 | 8290.08 |
| *Pseudalcantarea_macro* | night | -31.7 | C3 | 2696.56 | 1159.43 | 1305.762274 | 770.103 | 349.897 | 31.0207 | 331524 | 27640.7 | 89943.5 | 690556 | 224980 | 478.527 | 0 | 44.7028 | 1525.61 | 140973 | 305444 | 2940.36 |
| *Pseudalcantarea_macro* | day | -31.7 | C3 | 19607.5 | 3020.17 | 2951.495845 | 1835.96 | 590.526 | 58.9723 | 143986 | 80383 | 86986.3 | 2045337 | 535000 | 0 | 4331.77 | 119.936 | 201.825 | 169532 | 1135913 | 15417.8 |
| *Racinea_sinuosa** | night | -26.5 | C3 | 2679.2 | 2176.06 | 35326.83196 | 20170.3 | 51705.5 | 5694.33 | 81319.5 | 14509.8 | 69542.5 | 326134 | 48958.3 | 1496.8 | 2245.51 | 3990.39 | 12765.8 | 23796.4 | 36976.9 | 2983.28 |
| *Racinea_sinuosa** | day | -26.5 | C3 | 93256.4 | 122061 | 147193.2033 | 90564.2 | 70680.9 | 7423.87 | 255350 | 38830 | 57521.6 | 2375417 | 133437 | 61690.8 | 12757.3 | 11652.4 | 3390.19 | 78990.6 | 38920.5 | 12420.1 |
| *Tillandsia_demissa* | night | -22.0 | CAM | 173360 | 67712.3 | 553965.9783 | 371215 | 119133 | 13083.9 | 120183 | 76536 | 30313 | 1556260 | 136181 | 1664.62 | 0 | 6131.82 | 26687 | 31119.2 | 33969.2 | 5527.04 |
| *Tillandsia_demissa* | day | -22.0 | CAM | 158765 | 13986.1 | 1151450.623 | 860618 | 298516 | 46113.3 | 338139 | 89325.5 | 42707.7 | 626581 | 175276 | 858.026 | 18316.5 | 8212.1 | 31156.3 | 140640 | 58424.8 | 5418.42 |
| *Tillandsia_disticha* | night | -17.6 | CAM | 24425.3 | 182872 | 62074.05276 | 40982.1 | 58892.3 | 7539.57 | 86230.1 | 8485.76 | 47680 | 178037 | 259782 | 20588.9 | 2386.62 | 4272.59 | 121707 | 53301.3 | 38850.6 | 25165.6 |
| *Tillandsia_disticha* | day | -17.6 | CAM | 28046.7 | 197544 | 90973.96135 | 60379.3 | 69287.2 | 8525.77 | 112679 | 9379.57 | 66001.1 | 445008 | 402291 | 16007.1 | 2820.02 | 5472.46 | 42856.3 | 63124.2 | 29652.3 | 19970.5 |
| *Tillandsia_latifolia_aff_* | night | -15.2 | CAM | 48460.9 | 43563.1 | 948588.4 | 680268 | 298189 | 202638 | 105547 | 17019.1 | 93545 | 608340 | 631268 | 2019.5 | 2029.23 | 2826.78 | 177545 | 45227.8 | 39647.5 | 7845.46 |
| *Tillandsia_latifolia_aff_* | day | -15.2 | CAM | 76503.7 | 29098 | 767562.7 | 559016 | 438244 | 107900 | 115770 | 14401.1 | 20024.8 | 0 | 25245.7 | 0 | 11080.9 | 3935.76 | 77035.1 | 9139.19 | 270.396 | 7266.24 |
| *Tillandsia_ionantha* | night | -13.9 | CAM | 30064.5 | 212978 | 31849.72772 | 18958.6 | 26435 | 2899.85 | 132593 | 7230.12 | 18763.4 | 41869.8 | 55334.7 | 76799.8 | 0 | 2265.47 | 1331.71 | 20032.3 | 1582.92 | 17122.4 |
| *Tillandsia_ionantha* | day | -13.9 | CAM | 15077.2 | 69694.7 | 112151.2742 | 70267 | 67172.6 | 7425.4 | 97698.8 | 3958.2 | 17628.8 | 1824.07 | 10476.6 | 14966 | 0 | 1415.43 | 25590.6 | 11168.5 | 26.0446 | 3522.96 |
| *Tillandsia_juncea* | night | -15.35 | CAM | 192016 | 194108 | 4335.555556 | 2755.22 | 1933.79 | 324.13 | 209009 | 13192.8 | 37472.5 | 478669 | 222682 | 19780.3 | 7315.72 | 1481.47 | 8404.69 | 91140.3 | 7985.34 | 12816 |
| *Tillandsia_juncea* | day | -15.35 | CAM | 152129 | 212114 | 8213.951613 | 5005.51 | 7266.88 | 735.081 | 103364 | 5361.69 | 34776 | 986371 | 74566.2 | 62020.9 | 3114.14 | 262.341 | 16534.3 | 49801.4 | 8862.2 | 16189.3 |
| *Tillandsia_rauhii* | night | -20.7 | CAM | 10657.5 | 97889.1 | 152521.0644 | 102793 | 308752 | 62791.3 | 99131 | 585005 | 17994.1 | 0 | 9552.5 | 2313.12 | 5623.89 | 15596.5 | 97225.2 | 3154.88 | 23.1676 | 4747.5 |
| *Tillandsia_rauhii* | day | -20.7 | CAM | 3254.73 | 8743.77 | 430647.4064 | 312482 | 84098.9 | 133504 | 222864 | 355217 | 7071.44 | 0 | 1953.1 | 3776.55 | 6882.43 | 715.508 | 86852.8 | 5066.34 | 238.329 | 3108.98 |
| *Vriesea_longicaulis** | night | -26.1 | C3 | 22727.5 | 2417.06 | 65395 | 42303.9 | 121988 | 16435.3 | 167993 | 241416 | 44332.6 | 34393.3 | 62874.7 | 0 | 15843 | 21818.8 | 59858.9 | 91424.4 | 9614.67 | 424.667 |
| *Vriesea_longicaulis** | day | -26.1 | C3 | 11975.5 | 2080.19 | 136553.7864 | 91848 | 53220.5 | 6221.33 | 425287 | 70523 | 42437.8 | 150738 | 29580.3 | 631.286 | 14430 | 6544.78 | 17659.6 | 71504.3 | 40934.7 | 4216.75 |
| *Wallisia_lindeniana* | night | -26.9 | C3 | 97749 | 4325.73 | 19261.91099 | 11472.6 | 23731.3 | 2401.23 | 59984 | 520.052 | 19441.3 | 29373.3 | 19310.2 | 2737.41 | 101.859 | 7101.73 | 83284.7 | 24305.4 | 1559.29 | 4570.94 |
| *Wallisia_lindeniana* | day | -26.9 | C3 | 86534.2 | 1489.18 | 9321.625 | 5314.53 | 14333.3 | 1569.08 | 106834 | 812.975 | 10666.9 | 45074.6 | 17905.6 | 2156.68 | 2014.93 | 4916.85 | 67445.3 | 36604.1 | 1997.58 | 3057.43 |
| *(Tillandsia adpressifolia)* | na | -29 | na | na | na | na | na | na | na | na | na | na | na | na | na | na | na | na | na | na | na |
| *(Tillandsia curvispica)* | na | -22.9 | na | na | na | na | na | na | na | na | na | na | na | na | na | na | na | na | na | na | na |
| *(Tillandsia traunerii)* | na | -25.7 | na | na | na | na | na | na | na | na | na | na | na | na | na | na | na | na | na | na | na |

**SI_Table_2:** Abundance data for compounds identified by targeted metabolite analysis (4/6)

| species | sampling_time | isotope | PS | Glycine_2TMS | Glycine_3TMS | Isoleucine | Lactic_acid_DL | Leucine_1TMS | Leucine_2TMS | Phenylalanine | Proline | Putrescine | Serine | Succinic_acid | Threonic_acid | Threonine | Valine | phylogeny_label |
|---|---|---|---|---|---|---|---|---|---|---|---|---|---|---|---|---|---|---|
| *Alcantarea_regina** | day | -29.7 | C3 | 637.899 | 24956.7 | 9943.96 | 11070.1 | 16664.8 | 1501.14 | 2882.05 | 10363.7 | 6380.41 | 97510.7 | 2833.43 | 60208.5 | 35504.2 | 17165.4 | Atre_504a(=A. trepid |
| *Alcantarea_regina** | night | -29.7 | C3 | 5854.61 | 19429.3 | 5876.37 | 92.8884 | 4000.43 | 1636.7 | 9216.32 | 2467.62 | 60025 | 15210.5 | 18895.1 | 231319 | 8175.96 | 12256.2 | Atre_504a(=A. trepid |
| *Barfussia_laxissima* | night | -26.2 | C3 | 11889.4 | 8028.48 | 1453.56 | 59635.2 | 18138.8 | 0 | 537.838 | 9391.26 | 2834.44 | 72771.9 | 7328.19 | 3007.03 | 9437.01 | 2561.5 | Blax_B294 |
| *Barfussia_laxissima* | day | -26.2 | C3 | 10108.1 | 5059.1 | 1575.46 | 20347.2 | 13913 | 0 | 0 | 18055.1 | 4377.72 | 95822.6 | 1960.72 | 889.257 | 9980.29 | 2446.95 | Blax_B294 |
| *Tillandsia_australis* | night | -29.0 | C3 | 5638.63 | 82936.5 | 4497.24 | 46460.3 | 12715.1 | 2098.78 | 16949.6 | 35818.7 | 9867.02 | 232443 | 8215.73 | 13313.2 | 89927.9 | 15618.7 | Brom1_13 |
| *Tillandsia_australis* | day | -29.0 | C3 | 1133.23 | 10946.1 | 1381.41 | 47135.1 | 5916.72 | 840.502 | 1550.12 | 6132.46 | 4868.95 | 23336.4 | 5643.73 | 291447 | 6491.6 | 4698.59 | Brom1_13 |
| *Tillandsia_leiboldiana* | night | -31.3 | C3 | 0 | 8962.05 | 2118.11 | 39124.2 | 22270.6 | 1701.11 | 475.751 | 4321.68 | 173995 | 21151 | 2557.93 | 1185.88 | 4931.32 | 4703.39 | garden13 |
| *Tillandsia_leiboldiana* | day | -31.3 | C3 | 652.105 | 20196 | 2333.35 | 78299.5 | 12628.3 | 2038.21 | 853.421 | 6119.09 | 3322.2 | 18000.9 | 14716 | 19789 | 7597.94 | 5223.04 | garden13 |
| *Tillandsia_propagulifera* | night | -20.4 | CAM | 3979.49 | 83973.8 | 16561.6 | 9892.53 | 30866.5 | 6273.96 | 17840.3 | 27065 | 6467.11 | 30015.1 | 8772.66 | 2976.62 | 25237.4 | 31070.5 | garden20 |
| *Tillandsia_propagulifera* | day | -20.4 | CAM | 8163.17 | 191297 | 10805.5 | 242241 | 2551.56 | 4048.5 | 29444.4 | 8503.56 | 19711 | 25948.5 | 8344.35 | 5763.22 | 22716.5 | 20124.1 | garden20 |
| *Tillandsia_floribunda* | night | -18.6 | CAM | 28333.4 | 290642 | 11160.5 | 362613 | 37164.6 | 9908.32 | 5320.92 | 45993.3 | 12787.5 | 128153 | 9872.13 | 6908.7 | 55988.3 | 28582.3 | Brom1_211 |
| *Tillandsia_floribunda* | day | -18.6 | CAM | 13348.9 | 167992 | 2490.38 | 17431.7 | 45095.5 | 1125.19 | 1970.43 | 38592.6 | 14528.3 | 76393.4 | 23690.5 | 7897.36 | 43237.9 | 13401.4 | Brom1_211 |
| *Tillandsia_latifolia_ssp_latifolia* | night | -18.1 | CAM | 3140.95 | 16141.2 | 1655.4 | 63697.6 | 24704.2 | 2016.98 | 783.069 | 4013.15 | 2022.63 | 24058.4 | 3035.5 | 2852.46 | 5033.3 | 4507.14 | garden25 |
| *Tillandsia_latifolia_ssp_latifolia* | day | -18.1 | CAM | 761.631 | 16160.2 | 429.904 | 16737.5 | 8531.53 | 366.091 | 0 | 1833.36 | 4440.12 | 7837.79 | 29968.2 | 39072.7 | 2671.68 | 1391.97 | garden25 |
| *Vriesea_hitchcockiana* | night | -17.32 | CAM | 4314.69 | 70837.1 | 4871.01 | 39147.2 | 25654.4 | 2326.27 | 3507.27 | 53002.2 | 4157.49 | 104163 | 7880.74 | 2604.3 | 104130 | 14693.5 | garden39 |
| *Vriesea_hitchcockiana* | day | -17.32 | CAM | 1340 | 28610.1 | 905.668 | 9195.11 | 18477.8 | 487.003 | 862.343 | 29419.6 | 4026.75 | 66917.7 | 7467.18 | 1243.02 | 42632 | 6214.48 | garden39 |
| *Tillandsia_sphaerocephala°* | night | -15.7 | CAM | 15301.4 | 185457 | 4268.08 | 63012.4 | 27203.8 | 2517.66 | 4527.26 | 175879 | 24129.2 | 80479 | 6364.68 | 2094.13 | 53428.7 | 17080.8 | Brom1_11 |
| *Tillandsia_sphaerocephala°* | day | -15.7 | CAM | 1585.28 | 19724.8 | 1268.67 | 12090.6 | 18313.4 | 335.667 | 1551.92 | 35435.2 | 9088.81 | 16498.5 | 5438.36 | 1593.69 | 8933.69 | 6780.31 | Brom1_11 |
| *Tillandsia_somnians* | night | -26.9 | C3 | 1702.32 | 52162.6 | 4167.58 | 63240.2 | 20870.1 | 3142.54 | 1936.2 | 15167.1 | 14595.5 | 248025 | 4583.49 | 4354.4 | 35843.2 | 22254.1 | garden70 |
| *Tillandsia_somnians* | day | -26.9 | C3 | 4072.26 | 10994.3 | 3192.21 | 34474.2 | 12282.9 | 1551.74 | 777.189 | 3698.76 | 492594 | 19246.5 | 2735.77 | 5079.43 | 12624.8 | 8403.71 | garden70 |
| *Tillandsia_complanata* | night | -30.9 | C3 | 2889.88 | 13430.6 | 2725.52 | 33277 | 10250.2 | 822.645 | 2331.45 | 7625.76 | 3770.44 | 35821.6 | 10266.3 | 0 | 12218.1 | 6996.69 | garden90 |

**SI_Table_2:** Abundance data for compounds identified by targeted metabolite analysis (5/6)

| species | sampling_time | isotope | PS | Glycine_2TMS | Glycine_3TMS | Isoleucine | Lactic_acid_DL | Leucine_1TMS | Leucine_2TMS | Phenylalanine | Proline | Putrescine | Serine | Succinic_acid | Threonic_acid | Threonine | Valine | phylogeny_label |
|---|---|---|---|---|---|---|---|---|---|---|---|---|---|---|---|---|---|---|
| *Tillandsia_complanata* | day | -30.9 | C3 | 2616.48 | 122920 | 2878.95 | 95270.4 | 7263.72 | 1741.72 | 3482.92 | 58748.1 | 2702.22 | 92735.1 | 17795.3 | 386.534 | 17557.9 | 7596.36 | garden90 |
| *Lemeltonia_dodsonii* | night | -24.8 | C3 | 2657.44 | 44640.6 | 2686.73 | 28992 | 30687.5 | 3775.42 | 5624.99 | 16394.7 | 10725.9 | 173104 | 5573.95 | 19545.9 | 29308.4 | 10448.8 | Ldod_B127 |
| *Lemeltonia_dodsonii* | day | -24.8 | C3 | 9037.89 | 157514 | 5751.62 | 26232 | 10482.1 | 3849.44 | 5646.73 | 40497.7 | 6479.44 | 314787 | 8816.83 | 1940.08 | 156914 | 14844.9 | Ldod_B127 |
| *Tillandsia_striaca* | night | -15.4 | CAM | 3807.42 | 7205.32 | 1295.25 | 12526.9 | 20442.3 | 404.391 | 1027.23 | 8841.36 | 10647.5 | 12044.3 | 16628.7 | 10173 | 10276.5 | 6029.86 | TMM698_B |
| *Tillandsia_striaca* | day | -15.4 | CAM | 247.52 | 5916.63 | 1070.45 | 12710.5 | 17836 | 549.356 | 833.663 | 6397 | 10789.6 | 8461.04 | 15166.1 | 8372.62 | 8644.73 | 5477.18 | TMM698_B |
| *Tillandsia_arcuans(=syn.T.lajensis)** | night | -25.7 | C3 | 3179.67 | 8224.38 | 1544.41 | 54587.4 | 19604.5 | 1462.59 | 0 | 3359.24 | 2175.67 | 8271.45 | 12047.1 | 1968.38 | 2727.15 | 4205.87 | garden74 (=T.stenou |
| *Tillandsia_arcuans(=syn.T.lajensis)** | day | -25.7 | C3 | 2598.1 | 6572.39 | 971.346 | 19870.3 | 25277 | 328.599 | 0 | 2823.27 | 1528.85 | 4785.58 | 13627.3 | 2247.97 | 1826.02 | 2831.73 | garden74 (=T.stenou |
| *Tillandsia_australis°* | night | -26.1 | C3 | 6649.71 | 83754.3 | 29151.3 | 40179.1 | 0 | 5397.43 | 11596.1 | 24996 | 3101.64 | 346382 | 6425.87 | 9347.79 | 96638.9 | 73485.4 | Brom1_26 |
| *Tillandsia_australis* | night | -27.6 | C3 | 8184.57 | 150260 | 12882.6 | 29756.6 | 15339.9 | 2740.69 | 10579.4 | 25183.8 | 4389.52 | 494918 | 11003.9 | 36354.1 | 144556 | 34518 | Brom1_14 |
| *Tillandsia_australis°* | day | -26.1 | C3 | 13368.6 | 138994 | 34071.3 | 69039.3 | 0 | 7078.02 | 12709.4 | 38457.5 | 22362.4 | 612988 | 5334.06 | 6710.05 | 131537 | 91658.3 | Brom1_26 |
| *Tillandsia_australis* | day | -27.6 | C3 | 10084.7 | 177595 | 19306.9 | 79568.6 | 0 | 3531.98 | 11161 | 29302.3 | 9166.24 | 742807 | 5150.73 | 29508.2 | 234677 | 65346.4 | Brom1_14 |
| *Tillandsia_capillaris* | night | -15.0 | CAM | 1201.71 | 21847 | 2697.11 | 41493.8 | 21176.8 | 1287.48 | 3118.22 | 11924.4 | 9213.65 | 22360.6 | 4903.18 | 3133.36 | 8682.34 | 13298.1 | na |
| *Tillandsia_capillaris* | day | -15.0 | CAM | 481.984 | 12329.2 | 1078.15 | 15470.8 | 25522 | 581.384 | 1145.22 | 3638.59 | 69022.6 | 15020.5 | 8996.58 | 2423.34 | 4242.06 | 7626.55 | na |
| *Tillandsia_fasciculata°* | night | -16.1 | CAM | 3212.39 | 70858.8 | 9747.77 | 36496.5 | 18864.2 | 3677.46 | 3460.68 | 28118.5 | 18357.1 | 239328 | 8870.38 | 6155.38 | 105681 | 32859.4 | Brom1_29 |
| *Tillandsia_fasciculata* | night | -18.3 | CAM | 1512.96 | 79324 | 1994.82 | 19942.4 | 31597.9 | 1566.47 | 1872.78 | 26261.7 | 14599.6 | 82238.6 | 5382.61 | 82606.3 | 26432.6 | 7964.8 | Brom1_15 |
| *Tillandsia_fasciculata* | night | -14.5 | CAM | 0 | 6698.22 | 762.145 | 18884.6 | 0 | 583.733 | 0 | 1149.25 | 1018.05 | 21168.5 | 8446.02 | 64782.8 | 4811.64 | 2494.18 | Brom1_28 |
| *Tillandsia_fasciculata°* | day | -16.1 | CAM | 4824.4 | 70239.3 | 1371.11 | 12131.9 | 16715.4 | 496.923 | 1411.78 | 5803.24 | 6062.31 | 68246.4 | 6962.57 | 2005.17 | 16754.9 | 8012.76 | Brom1_29 |
| *Tillandsia_fasciculata* | day | -18.3 | CAM | 18817.2 | 90644.3 | 13886.7 | 89475.5 | 25483.5 | 4276.59 | 12411 | 84263.4 | 64028.5 | 295967 | 34343.9 | 48658 | 137129 | 35166.4 | Brom1_15 |
| *Tillandsia_fasciculata* | day | -14.5 | CAM | 1411.96 | 26211.3 | 2785.94 | 23712.9 | 0 | 1069.21 | 1881.94 | 12474.5 | 43238.3 | 37440.2 | 13731.5 | 4404.18 | 16356.7 | 7300.41 | Brom1_28 |
| *Tillandsia_floribunda* | night | -20.3 | CAM | 16851.7 | 182738 | 7405.18 | 258034 | 26501.5 | 7092.93 | 4497.15 | 48490.3 | 5141.97 | 88720.7 | 16343.9 | 10967.4 | 40186.8 | 19651.5 | Brom2_16 |
| *Tillandsia_floribunda* | night | -18.9 | CAM | 37608.7 | 444922 | 5203.85 | 69298.8 | 22420.3 | 2816.88 | 6663.56 | 36609.9 | 9588.07 | 230870 | 14113.9 | 11768.7 | 106405 | 24868.5 | Brom1_16 |
| *Tillandsia_floribunda* | day | -20.3 | CAM | 9015.95 | 80028.9 | 1301.08 | 36335.5 | 55930.2 | 761.132 | 1227.55 | 37524.6 | 2135.21 | 40232.5 | 24260.3 | 6274.39 | 24612.2 | 5513.68 | Brom2_16 |
| *Tillandsia_floribunda* | day | -18.9 | CAM | 32058.1 | 574774 | 3640.54 | 12410.3 | 47092.6 | 1076.76 | 3860.93 | 41631.1 | 27560.5 | 193138 | 39038.7 | 11552.5 | 105021 | 24639.6 | Brom1_16 |
| *Tillandsia_sphaerocephala* | night | -16.2 | CAM | 2607.38 | 34359.9 | 0 | 36966.8 | 28607.2 | 883.281 | 2000.34 | 23591.3 | 42476.8 | 15937.7 | 12497 | 4534.57 | 14437.7 | 8023.67 | Brom1_12 |
| *Tillandsia_sphaerocephala* | night | -15.7 | CAM | 5775.53 | 98365.2 | 1895.53 | 22652.7 | 34420.9 | 605.452 | 798.243 | 26354.2 | 9943.82 | 28439.1 | 5410.31 | 2364.63 | 19183.3 | 10411 | Brom1_23 |

**SI_Table_2:** Abundance data for compounds identified by targeted metabolite analysis (6/6)

| species | sampling_time | isotope | PS | Glycine_2TMS | Glycine_3TMS | Isoleucine | Lactic_acid_DL | Leucine_1TMS | Leucine_2TMS | Phenylalanine | Proline | Putrescine | Serine | Succinic_acid | Threonic_acid | Threonine | Valine | phylogeny_label |
|---|---|---|---|---|---|---|---|---|---|---|---|---|---|---|---|---|---|---|
| *Tillandsia_sphaerocephala* | day | -16.2 | CAM | 1557.57 | 22945 | 0 | 19551 | 53944.1 | 183.133 | 1135.25 | 18919.1 | 1294.39 | 6509.67 | 40949 | 5655.14 | 7962.65 | 7659.45 | Brom1_12 |
| *Tillandsia_sphaerocephala* | day | -15.7 | CAM | 1585.85 | 12317.1 | 826.415 | 15871.1 | 16852.3 | 368.005 | 74.8032 | 12001.1 | 2092.88 | 4695.71 | 21969.4 | 2616.82 | 3045.55 | 4480.43 | Brom1_23 |
| *Pseudalcantarea_macropetala* | night | -31.7 | C3 | 6234.81 | 13713.9 | 3453.02 | 11457.9 | 10627.2 | 830.801 | 11431.4 | 12817 | 17266.9 | 231171 | 1028.42 | 0 | 26805.9 | 7339.84 | Pmac_B742 |
| *Pseudalcantarea_macropetala* | day | -31.7 | C3 | 10931.4 | 24991.3 | 2272.96 | 53140.9 | 13154.1 | 933.767 | 2786.37 | 18742.7 | 11268.4 | 104222 | 4909.53 | 6332.47 | 26716.1 | 8214.63 | Pmac_B742 |
| *Racinea_sinuosa** | night | -26.5 | C3 | 2616.69 | 36569.9 | 2854.1 | 45073.6 | 33200.9 | 1702.53 | 1568.51 | 7329.7 | 42503.9 | 27244 | 9700.69 | 4974.44 | 9685.81 | 8730.8 | Rspi_B1596 |
| *Racinea_sinuosa** | day | -26.5 | C3 | 947.521 | 28446.9 | 2521.62 | 12362.3 | 47814.6 | 1013.84 | 2132.56 | 12811.3 | 49876.4 | 33472.4 | 13917.1 | 4771.11 | 15000.4 | 7149.55 | Rspi_B1596 |
| *Tillandsia_demissa* | night | -22.0 | CAM | 951.63 | 26315.3 | 1041.44 | 27057.6 | 31446.4 | 786.005 | 1454.78 | 19790.4 | 40105.2 | 13808.5 | 7477.12 | 2798.18 | 11746.4 | 6355.65 | Tdem_B1373 |
| *Tillandsia_demissa* | day | -22.0 | CAM | 510.182 | 31884.3 | 955.74 | 13370.6 | 17762.1 | 528.623 | 2006.34 | 12940.3 | 191933 | 15735.8 | 57388 | 37237.6 | 18414.7 | 2136.44 | Tdem_B1373 |
| *Tillandsia_disticha* | night | -17.6 | CAM | 2614.72 | 48607.9 | 935.875 | 25327.5 | 22006.7 | 866.906 | 552.398 | 7393.43 | 13565.3 | 30644 | 10209.5 | 28707 | 13794.7 | 6829.69 | Tdis_B1595 |
| *Tillandsia_disticha* | day | -17.6 | CAM | 9930.72 | 113600 | 1958.74 | 19744.1 | 19096.3 | 949.396 | 1215.89 | 8957.97 | 11273.4 | 42773.5 | 9145.31 | 9864.13 | 22608.5 | 10633.3 | Tdis_B1595 |
| *Tillandsia_latifolia_aff_divaricata* | night | -15.2 | CAM | 3183.12 | 66985.9 | 10707 | 139150 | 20208.4 | 8753.62 | 8363.17 | 23600.3 | 11431.6 | 74872.7 | 6347.73 | 3964.14 | 24102 | 22702.6 | Tdiv_B1594 |
| *Tillandsia_latifolia_aff_divaricata* | day | -15.2 | CAM | 1406.24 | 24753.9 | 690.815 | 16024.4 | 10317.2 | 498.202 | 0 | 6793.88 | 3092.53 | 11967.5 | 9169.66 | 7691.54 | 4138.15 | 2642.95 | Tdiv_B1594 |
| *Tillandsia_ionantha* | night | -13.9 | CAM | 1451.24 | 26638.1 | 1528.54 | 24571.2 | 33174.9 | 917.005 | 415.693 | 34083.1 | 144360 | 21213.7 | 5537.3 | 1122.7 | 5049.65 | 7497.03 | Tion_B84 |
| *Tillandsia_ionantha* | day | -13.9 | CAM | 1027.01 | 18525 | 1131.99 | 31366.9 | 25061.7 | 557.729 | 172.922 | 20846.8 | 49479.4 | 10474 | 8056.09 | 577.479 | 2945.98 | 4791.47 | Tion_B84 |
| *Tillandsia_juncea* | night | -15.35 | CAM | 2878.14 | 50696.9 | 2305.1 | 60031.5 | 24590.7 | 2526.5 | 1395.6 | 16839.9 | 75097 | 65941 | 5205.02 | 976.498 | 16310.8 | 12226.5 | Tjunc_633 |
| *Tillandsia_juncea* | day | -15.35 | CAM | 3069.46 | 54006.6 | 1076.24 | 18900.4 | 29187.8 | 498.387 | 526.371 | 5985.35 | 26915.8 | 37525.8 | 3676.99 | 1440.62 | 9089.11 | 7945.43 | Tjunc_633 |
| *Tillandsia_rauhii* | night | -20.7 | CAM | 0 | 3797 | 412.995 | 14048.9 | 23203.9 | 440.025 | 0 | 865.99 | 4266.41 | 3069.11 | 7212.3 | 15686.2 | 953.713 | 1429.75 | Trau_B92 |
| *Tillandsia_rauhii* | day | -20.7 | CAM | 1780.64 | 4942.11 | 0 | 0 | 44940.2 | 0 | 0 | 0 | 48492.2 | 380.027 | 22978.5 | 31422.8 | 0 | 884.706 | Trau_B92 |
| *Vriesea_longicaulis** | night | -26.1 | C3 | 3101.33 | 9693.19 | 5388.64 | 38010.7 | 19162.3 | 4090.25 | 2871.31 | 23101.7 | 186020 | 53814.7 | 10495.1 | 1508.11 | 21170.5 | 23032.4 | Vitat_574 |
| *Vriesea_longicaulis** | day | -26.1 | C3 | 6675.39 | 15858.5 | 0 | 24485.8 | 5855.68 | 689.393 | 668.859 | 3227.77 | 55642.7 | 16141.6 | 5521.87 | 3332.79 | 6033.57 | 3453.98 | Vitat_754 |
| *Wallisia_lindeniana* | night | -26.9 | C3 | 795.262 | 10322.6 | 3291.78 | 40908.2 | 56069.1 | 1467.72 | 1994.97 | 9513.9 | 8295.6 | 18994.6 | 6962.07 | 7005.97 | 8368.66 | 11987.3 | Wlin_B758 |
| *Wallisia_lindeniana* | day | -26.9 | C3 | 298.3 | 4906.48 | 1166.58 | 14610.4 | 45032.2 | 608.075 | 815.675 | 7701.1 | 8416.48 | 16437.6 | 7815.6 | 9590.78 | 4856.6 | 6475.43 | Wlin_B758 |
| *(Tillandsia adpressifolia)* | na | -29 | na | na | na | na | na | na | na | na | na | na | na | na | na | na | na | Brom_Garden50 |
| *(Tillandsia curvispica)* | na | -22.9 | na | na | na | na | na | na | na | na | na | na | na | na | na | na | na | Tcur_B1597 |
| *(Tillandsia traunerii)* | na | -25.7 | na | na | na | na | na | na | na | na | na | na | na | na | na | na | na | Brom_Garden36 |

#### SI_Table_3: Genes putatively affected by adaptive protein evolution as revealed by branch-specific tests for selection.

| Gene | Test 3 | | | | Test 4 day | | | Test4 night | | | Description | References |
|---|---|---|---|---|---|---|---|---|---|---|---|---|
| | Tspha | Taust | Tfasc | Tflor | Tspha | Tfasc | Tflor | Tspha | Tfasc | Tflor | | |
| Aco020962† | - | - | ** | - | ** | ** | ns | ** | ** | ns | Enolase EC4.2.1.11, phosphoenolpyruvate hydratase. Last step of glycolysis which catalyses the reversible conversion of 2-phosphoglycerate into phosphoenolpyruvate (PEPC). PEPC is a key-enzyme in C4 and CAM photosynthesis | PEPC evolution: Christin et al. 2014 Wai et al. 2017 |
| Aco012463‡ | - | - | - | - | ns | ns | ns | ns | ns | ns | E3SUMO: E3-SUMO protein ligase SIZ1 controls phosphate deficiency response and regulates drought response | Miura et al. 2005 Catala et al. 2007 |
| Aco005505‡ | - | - | ns | - | ns | ** | ns | ns | ns | ns | LACS or ACSL, also called FATP: Long chain fatty acid CoA ligase. In plants, linked to cuticle growth (wax content), thus conferring susceptibility to drought stress. | Pulsifer 2012 Weng et al. 2010 |
| Aco009511‡ | - | - | - | - | ns | ns | ns | ns | ns | ns | ERF: Ethylene responsive transcription factor. TF present in numerous adaptive responses to a wide variety of biotic and abiotic stresses. In Pepper (*Capsicum* sp.), enhances drought tolerance. | Fujimoto et al. 2000 Hong et al. 2017 |
| Aco005399‡ | - | - | - | - | ns | ns | ns | ns | ns | ns | Protein of unknown function DUF1645, in rice this protein domain confers drought resistance | Cui et al. 2016 |
| Aco000648(‡) | - | - | - | - | ns | ns | ns | ns | ns | ns | Chaperone protein DnaJ: stress-induced heat shock protein (Hsp40), co-chaperone of Hsp70, large family of heat-shock proteins. Hsp plastid concentration promotes thermotolerance in C3, C4 and CAM species. | |
| Aco018002(‡) | - | - | - | - | ns | ns | ns | ns | ns | ns | PARG2: Poly-ADP ribose glycohydrolase 2, response to DNA damage, and immune response. PARG1 is involved in drought stress response. PARG1, PARG2, PARP1 and PARP2 are suggested to be linked by regulatory interactions. | Song et al. 2015 Li et al. 2011 |

| | | | | | | | | | | | | |
|---|---|---|---|---|---|---|---|---|---|---|---|---|
| Aco003224(‡) | - | - | ns | - | ns | ns | ns | ns | ** | ns | MYB: DNA-binding protein MYB transcription factor. Can be involved in many biotic and abiotic stress responses. | Ambawat et al. 2013 |
| Aco007014(‡) | - | - | - | - | ** | ** | ** | ** | ** | ** | LPIN3: Phosphatidate phosphatase, key regulatory enzyme in fatty acid metabolism. Potentially involved in heat-stress and drought-stress response. | Carman 2009<br>De Bigault Du Granrut, & Cacas 2016 |
| Aco013659 | ** | ** | ** | - | ** | ** | ns | ** | ns | * | SAM Mtase: S-adenosyl-L-methionine synthase 2516 (c.f. gene above), enzyme of the cysteine and methionine metabolism pathways (KEGG EC2.5.1.6, ec00270) | Joshi & Chiang 1998<br>Roje 2006 |
| Aco003279 | - | ns | - | - | ns | ns | ** | ns | ns | ** | SAM: S-adenosyl-L-methionine dependent methyltransferase superfamily protein. The methyl group donor has a regulatory function in many metabolic pathways. | Joshi & Chiang 1998<br>Roje 2006 |
| Aco002187 | - | - | - | - | ns | ns | ns | ns | ** | ns | TUBCGP: Gamma-tubulin complex component, highly conserved | Kong et al. 2010 |
| Aco012435 | - | - | - | - | ns | ns | ns | ns | ns | ns | G6PD: Glucose6phosphate dehydrogenase, first enzyme of the pentose-phosphate pathway. | Hauschild & von Schaewen 2003 |
| Aco001521 | - | - | - | - | ns | ns | ns | ns | ns | ns | CECR2 : Cat eye syndrome critical region protein-like protein | |
| Aco008752 | - | - | - | - | ns | ns | ns | ns | * | ns | Nucleoporin-related, highly conserved, few studies in plants | Tamura et al. 2010 |
| Aco011527 | - | - | - | - | ns | ** | ns | ns | ns | ns | CLUH: Clustered mitochondria protein-like protein, regulates mitochondrial biogenesis by binding mRNAs of nuclear-encoded mitochondrial proteins | Gao et al. 2014 |
| Aco013210 | - | - | ns | - | ns | ns | ns | ns | * | ns | PPR: Pentatricopeptide repeat containing protein, one of the largest protein families in plants, involved in many different biological processes. | Zhang et al. 2017<br>Barkan & Small 2014 |
| Aco011299 | - | - | - | - | ns | ns | ns | ns | ns | ns | PPR: Pentatricopeptide repeat containing protein, one of the largest protein families in plants, involved in many different biological processes. | Zhang et al. 2017<br>Barkan & Small 2014 |

| | | | | | | | | | | | |
|---|---|---|---|---|---|---|---|---|---|---|---|
| Aco007782 | - | - | - | - | ns | ns | ns | ns | ns | ns | InvB: Plant neutral invertase family protein Active in plant biotic stress response involving sucrose | Tauzin & Giardina 2004 |
| Aco008802 | - | - | - | - | ns | ns | ns | ns | ns | ns | FBT: Folate biopterin (vitamin B9) transporter, promoting biosynthesis of amino acids such as methionine, serine, histidine | Bedhomme et al. 2005 |
| Aco007483 | - | - | - | - | - | - | - | - | - | - | CAS1: Cyclo-artenol synthase1, cycloartenol, important precursor of sterols in photosynthetic organisms | Babiychuk et al. 2008 |

† : possible link to CAM photosynthesis; ‡ : potentially related to general drought / heat stress response syndrome; (‡): indirectly linked to drought stress response; Differential expression analysis, FDR significance thresholds: ****** : < 1%,, ***** : <5%, **ns** : >5% ; - : not differentially expressed.
